# Unsupervised learning predicts human perception and misperception of gloss

**DOI:** 10.1101/2020.04.07.026120

**Authors:** Katherine R. Storrs, Barton L. Anderson, Roland W. Fleming

## Abstract

Reflectance, lighting, and geometry combine in complex ways to create images. How do we disentangle these to perceive individual properties, like surface glossiness? We suggest that brains disentangle properties by learning to model statistical structure in proximal images. To test this, we trained unsupervised generative neural networks on renderings of glossy surfaces and compared their representations with human gloss judgments. The networks spontaneously cluster images according to distal properties such as reflectance and illumination, despite receiving no explicit information about them. Intriguingly, the resulting representations also predict the specific patterns of ‘successes’ and ‘errors’ in human perception. Linearly decoding specular reflectance from the model’s internal code predicts human gloss perception better than ground truth, supervised networks, or control models, and predicts, on an image-by-image basis, illusions of gloss perception caused by interactions between material, shape, and lighting. Unsupervised learning may underlie many perceptual dimensions in vision, and beyond.

## Main

A photograph of a glass of water might consist of large bright patches, sparkling dots, and low-contrast blurs—yet we immediately see these as reflections on the water’s surface, bubbles within, and smudges on the glass. Identifying the distal physical causes of proximal image features is widely considered to be the central challenge of vision ^1–7^. Yet how we infer the outside world from ambiguous sensory data remains mysterious.

Somehow the visual system infers the combination of distal scene variables that most plausibly explains the proximal sensory input ^4, 8–11^. Yet, a major unsolved question is how the candidate explanations of proximal inputs came to be known in the first place. Our visual systems did not – and do not – have access to ground truth information about the number or kinds of distal sources of image structure that operate in the world. Any knowledge about the world must have been acquired from exposure to proximal stimuli over evolutionary and/or developmental time scales ^12–14^.

Here we explore the intriguing possibility that visual systems might be able to discover the operation of distal scene variables by learning statistical regularities in proximal images, rather than through learning an explicit mapping between proximal cues and known distal causes. Specifically, we show that by learning to efficiently compress and spatially predict images of surfaces, an unsupervised generative deep neural network (DNN) not only spontaneously clusters inputs by distal factors like material and illumination, but also strikingly reproduces many characteristic ‘misperceptions’ of human observers.

Two general principles motivate our approach. The first is that variability in proximal sensory stimuli is caused by variations in a lower-dimensional set of environmental factors (such as shape, reflectance, and illumination). This implies that the variation between images can be captured in a more compact or simple way when represented in terms of their underlying causes, rather than (say) in terms of pixels ^15–17^. A machine learning model encouraged to discover a compact representation of images might therefore converge on the finite sources that generate image variability. Identifying and disentangling these distal sources is only possible, however, if a second principle holds: different distal sources must generate statistically distinguishable effects in the proximal input ^18^. This seems intuitively true – for example, changes in illumination generate different kinds or patterns of variability in images than changes in surface material do. Based on these two principles, we reasoned that it should be possible for a sufficiently powerful statistical learning model to discover the existence of distal variables without *a priori* knowledge of either the number or kinds of distal variables that exist in the world, based solely on the variability they generate in images.

The idea that our perceptual systems exploit statistical regularities to derive information about the world has a long and venerable history in both psychology and neuroscience ^19–21^. For example, neural response properties in early visual cortex are well predicted by models trained to generate sparse codes for natural image patches ^19, 21–24^. Unfortunately, such ‘efficient coding’ approaches have not yet scaled beyond the initial encoding of images. One of the main motivations for the present work was to determine whether such ideas could provide leverage into mid-level scene understanding, i.e., inferring the distal physical causes of sense data.

Even if different distal factors have different statistical effects on images, and a sufficiently powerful unsupervised neural network is able to learn these, its success in disentangling different factors is unlikely to be perfect. That is, the network would sometimes misattribute the distal causes responsible for the data. However, we regard such misattributions as a potential strength; there are well documented examples where the human visual system systematically misattributes image structure to the wrong distal source—failures of ‘perceptual constancy’ ^1, 5, 25–33^. We were interested in whether the pattern of successful and unsuccessful attributions made by human observers would also be exhibited by networks that failed to fully disentangle distal scene variables. The goal was not to understand visual processes as an estimation of ground truth, but rather, to understand why our visual systems extract what they do about the world in the absence of access to ground truth knowledge.

One of the most striking patterns of successes and failures in estimating distal scene variables occurs in the perception of surface gloss ^3, 5, 31, 34–37^. Gloss perception is a paradigmatic case of a perceptual judgment where multiple physical effects must be separated. The pattern of specular reflections can change dramatically as a function of a surface’s 3D shape, illumination direction, and the observer’s viewpoint. Indeed, psychophysical evidence has shown that the perception of gloss in human observers depends not only on specular reflectance, as expected ^32, 38–44^, but also on lighting and shape ^33–37, 45^. We were interested in whether the specific pattern of these complex interactions could be a consequence of the visual system having learned to approximately disentangle distal sources from their effects on image structure.

Our work exploits DNN methods that have emerged for learning sophisticated models of image structure in the form of latent variables, which summarise how images differ from one another ^46–50^. DNNs achieve complex transformations by passing input data through layers of units that apply weighted summations and non-linearities, roughly mimicking the operations of biological neurons ^51–55^. During training, connections between units are iteratively adjusted to improve the network’s performance on some learning objective.

In supervised learning, networks are directly told what label to output for every input image in a training dataset, from which they learn to subsequently output appropriate labels for new test images. Supervised DNNs have revolutionised computer vision, achieving near-human object and face recognition ^56–59^, and are the best extant models of late ventral stream function ^51–53, 60–63^. However, unlike humans, they are often fragile to tiny image perturbations ^64, 65^ and over-rely on local texture ^66, 67^. As models of biological gloss perception, it is unclear from where the necessary training labels could come.

In unsupervised learning, training objectives encourage networks to learn statistical regularities in the training data without being given any explicit labels. For example, autoencoder networks are trained to compress training images into compact descriptions, then reconstruct them as accurately as possible ^50, 68, 69^. Here, we use a variant known as a PixelVAE ^47, 48^, which learns to both summarise and spatially predict images, in two connected processing streams (Figure 1C). It is a generative model that can create completely novel images with high-order statistical structure similar to the training data—in our case, images of glossy and matte surfaces.

**Figure 1:**
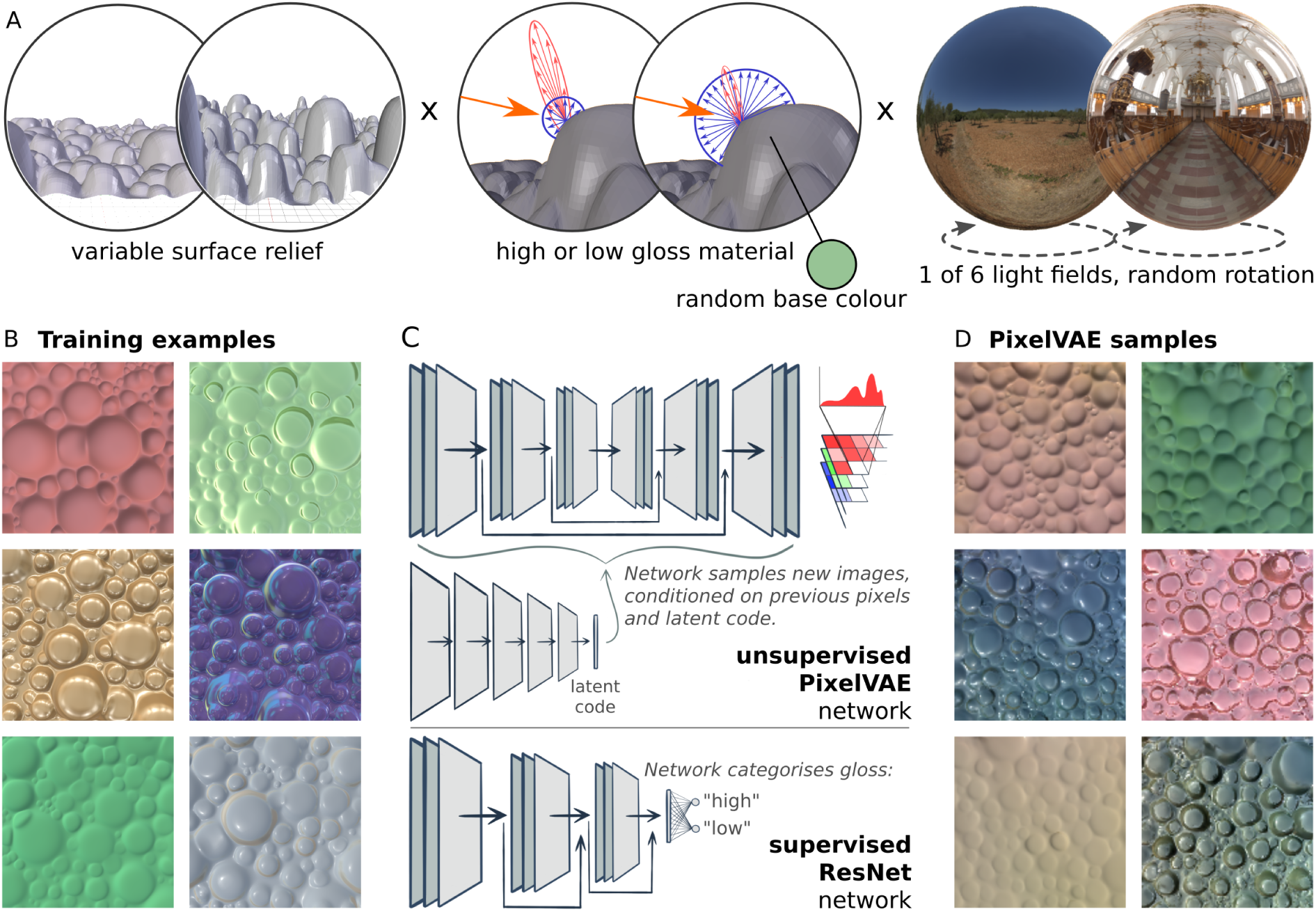
An unsupervised neural network learns to generate plausible novel images from a simulated visual world. (A) The simulated physical world consisted of a bumpy surface viewed from above. The scene varied in depth of surface relief (left), surface base colour and whether the surface material had high or low gloss (middle), and illumination environment (right). (B) Six examples from a training set of 10,000 images rendered from the simulated world by randomly varying the world factors. (C) The dataset was used to train an unsupervised PixelVAE network (upper panel). The network learns a probability distribution capturing image structure in the training dataset, which can be sampled from one pixel at a time to create novel images with similar structures. A sub-stream of the network learns a highly compressed 10-dimensional latent code capturing whole-image attributes, which is used to condition the sampling of pixels. The same dataset was used to train a supervised ResNet network, which output a classification of “high gloss” or “low gloss” for each input image. The penultimate layer of the supervised network is a 10-dimensional fully-connected layer, creating a representation of equivalent dimensionality for comparison to the unsupervised model’s latent code. (D) Six example images created by sampling pixels from a trained PixelVAE network, conditioning on different latent code values to achieve diverse surface appearances. These images are not reconstructions of any specific images from the training set, but are completely novel samples from the probability distribution the network has learned.

Our main finding is that the representation of gloss learned by an unsupervised PixelVAE network closely matches the pattern of successes and failures in perceived gloss shown by human observers. The unsupervised models better match human data than do a range of supervised networks and simpler comparison models, suggesting that the learning process by which different distal sources are disentangled may play a fundamental role in shaping our visual experience of the world, and providing a potential answer to the conundrum of how we learn to see without explicit training.

## Results

To test whether an unsupervised DNN can learn human-like gloss perception, we rendered 10,000 images from a virtual world consisting of frontal views of bumpy surfaces with either high (‘gloss’) or low (‘matte’) specular reflectance. Using renderings grants tight control over the statistics of the training environment, allowing us to guarantee that reflectance could not be trivially decoded from raw images and that physical factors varied independently of one another. Each image had a different random configuration of bumps, depth of surface relief, and colour, and was illuminated by one of six natural light fields (Figure 1A-B). We then trained ten different instances of a PixelVAE ^47, 48^ network with different initial random weights on this dataset, to ensure results were robust to representational differences between training instances of the same architecture ^70^. The network culminates in a probability distribution over pixel values (Figure 1C). Its training objective is to adjust the shape of this distribution in order to increase the likelihood of the training images under it, leading to a model of the structure and variability within and across images.

New images can be created from the unsupervised PixelVAE model. These are generated pixel-by-pixel from the top left corner by probabilistically sampling from the network’s learned distribution, conditioned both on previous pixels and values in the model’s 10-dimensional latent code (Figure 1C). The representations in this latent code are a highly compressed representation of whole-image properties, and are the focus of all subsequent analyses. After training, all ten instances of the model could generate wholly novel images that look like plausible surfaces (Figure 1D).

As a comparison DNN, we also trained ten instances of a supervised ResNet ^71^ network to classify the same images as high or low gloss, using ground truth (high or low specular reflectance in the rendering parameters) as training labels. Its mean classification accuracy was 99.4% +/-SD 0.001. This supervised model also contained a 10-dimensional fully-connected layer prior to its two-unit output classification layer, which we treated as its high-level latent code for comparisons with the unsupervised model (Figure 1C).

### An unsupervised generative model disentangles distal scene properties

We are interested in the extent to which models transform raw images into a feature space within which different physical causes are disentangled ^12, 14, 16, 72^. Surfaces with similar reflectance properties may occupy very disparate points in raw pixel space, but should cluster together in the feature space of a good perceptual model. Although the unsupervised PixelVAE model’s training objective deals only with proximal image data, after training on the rendered dataset, we found that distal scene properties—such as gloss and lighting—spontaneously clustered within the networks’ 10-dimensional latent codes (cf. ^47, 50, 73^). Visualising in two dimensions using tSNE ^74^ strikingly reveals that low-gloss images cluster together, while high-gloss images form multiple tight clusters, corresponding to different light fields (Figure 2A). Within each light-field cluster, images are arranged by the angle of illumination, as well as by surface relief, with flatter surfaces occupying nearby points, and bumpier surfaces more distant points. This shows that without explicit labels, the unsupervised model learns at least partially to reorganise stimuli by their physical properties, one of the core challenges of mid-level vision.

**Figure 2:**
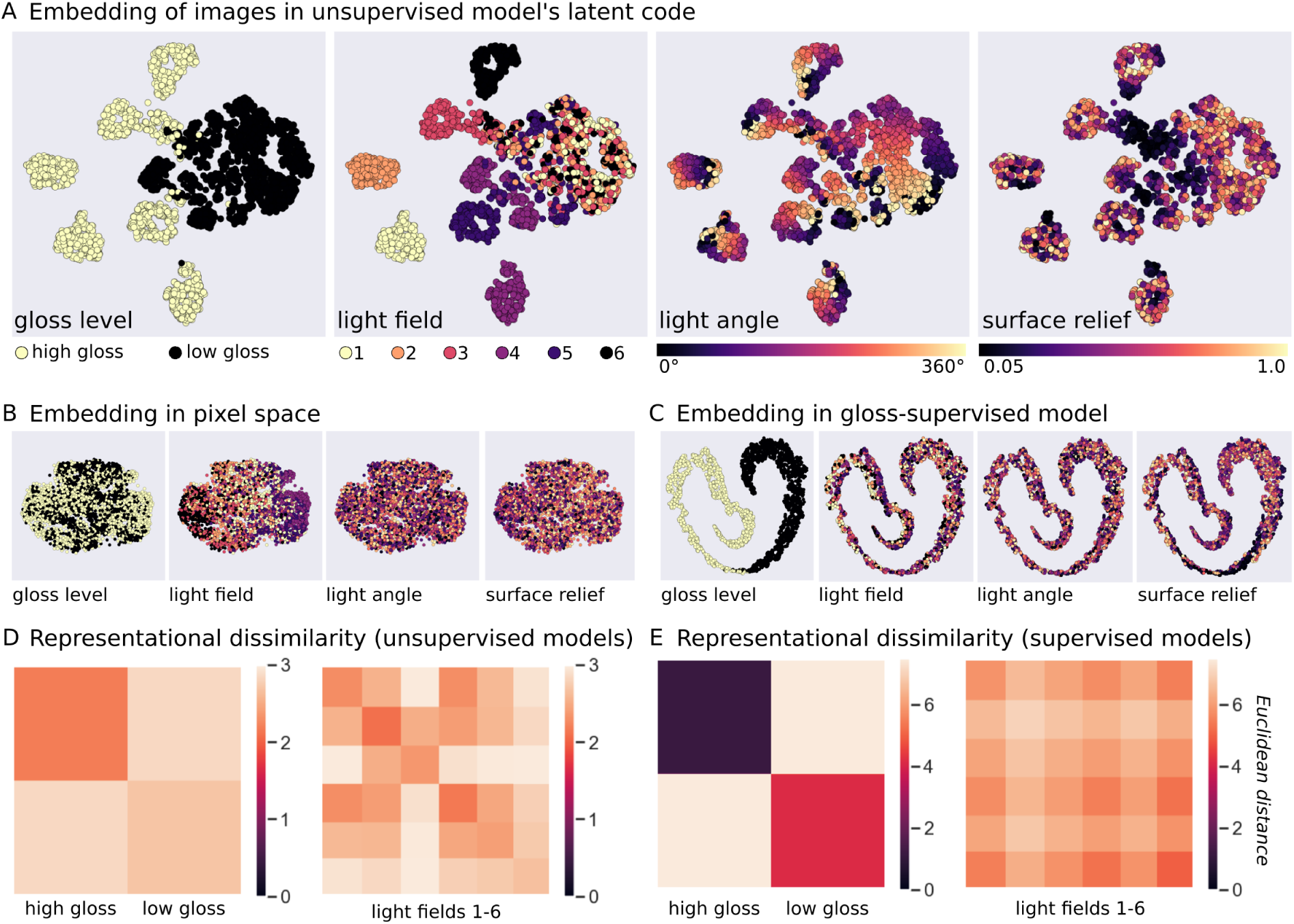
World factors are disentangled in the unsupervised model’s latent code. (A) Visualisations of distances between 4,000 images, in the 10D latent code of one unsupervised PixelVAE network, projected into two dimensions using tSNE. Unsupervised learning spontaneously disentangles the underlying world factors, arriving at a nested representation. Images of low-gloss surfaces form one large cluster while images of high-gloss surfaces form multiple small subclusters (left panel), according to the light field used to render the image (centre left panel). Lighting direction varies smoothly within each subcluster (centre right panel), and surfaces with low relief are closer to one another than are those with high relief (right panel). (B) tSNE visualisation of the same images in raw pixel space, where world factors are thoroughly entangled. (C) tSNE visualisation of the same images in the 10D latent code of one gloss-supervised ResNet network, showing a less rich representation. High- and low-gloss images form two clearly-separated clusters, but other world factors are entangled. (D) Dissimilarity matrices showing Euclidean distance between all pairs of images in the latent representations of all PixelVAE networks, averaged across surfaces with the same (left) gloss level, or (right) illumination. Pairs of images belonging to the same gloss/lighting condition (diagonal blocks in each matrix) were represented more similarly than pairs of images belonging to different conditions (off-diagonal blocks) (gloss: *t*_9_ = 16.73, *p* < 0.001, d = 0.97, 95% CI = [0.37-0.46]; light-field: *t*_9_ = 29.76, *p* < 0.001, d = 0.95, 95% CI = [0.36-0.41]). (E) Corresponding dissimilarity matrices calculated from the representations in the 10D latent codes of the supervised networks reveals stronger clustering by the task-relevant dimension of gloss, and weaker clustering by light-field (model×factor interaction *F_1,18_* = 9878.34, *p* < 0.001, η^2^ = 0.99).

The emergence of this clustering of images by scene properties is far from trivial. It was not caused by raw image similarities, since tSNE visualisation of the same images in raw pixel space showed a tight entangling of scene properties (Figure 2B). Other linear and non-linear pixel embeddings such as MDS and LLE ^75^ also failed to separate low-from high-gloss surfaces (Supplementary Figure 2). When the same visualisation was applied to the 10D layer of the gloss-supervised models, high and low gloss images were neatly separated, but other world factors were intermixed (Figure 2C). Similar qualitative patterns held for all ten instances of both unsupervised and supervised models.

To quantify these clustering effects, we used representational similarity analysis ^76^ (Figure 2D). The results support the tSNE visualisations. Pairs of images belonging to the same gloss condition (both glossy or both matte), corresponded to closer points in the unsupervised models’ 10D latent codes than pairs of images belonging to different gloss conditions (repeated-measures t-test comparing average distances between same-vs different-gloss image pairs, across network training instances: *t*_9_ = 16.73, *p* < 0.001, Cohen’s d = 0.97, 95% CI of difference = [0.37–0.46]). Likewise, pairs of images illuminated by the same light field had more similar latent representations than those lit by different light fields (*t*_9_ = 29.76, *p* < 0.001, d = 0.95, 95% CI = [0.36–0.41]). In the supervised models, clustering was dominated by gloss (Figure 2E; two-way mixed-effects ANOVA interaction between model type and scene factor *F_1,18_* = 9878.34, *p* < 0.001, η^2^ = 0.99; follow-up tests show far stronger gloss clustering in supervised than unsupervised models *t_18_* = 99.39, *p* < 0.001, d = 44.45, 95% CI = [6.42–6.66], but stronger light-field clustering in unsupervised models, *t_18_* = −19.90, *p* < 0.001, d = 8.90, 95% CI = [0.27–0.32]). Thus, while the supervised model optimizes disentanglement of the single physical property on which it is trained, the unsupervised model spontaneously discovers multiple scene factors contributing to image structure.

### The unsupervised model predicts human gloss perception for novel images

Our central question was whether the spontaneous separation of high and low gloss images in the unsupervised model could capture human gloss judgments. To derive quantitative gloss predictions from the models, a linear support vector machine (SVM) classifier was trained to find the hyperplane in the 10D latent code of each network that best separates high from low specular reflectance images (Figure 3A). Although this evaluation step involves label-based decoding, it is simply a formal way of quantifying the degree and form of the disentanglement. In neuroscience, information that is available directly via a linear readout from units or neurons is generally considered to be explicitly represented by a model or brain region (e.g. ^16, 77–81^). The linear classifier does not provide the model with any new information, but merely measures the relative placement of different classes of images within its existing feature space.

**Figure 3:**
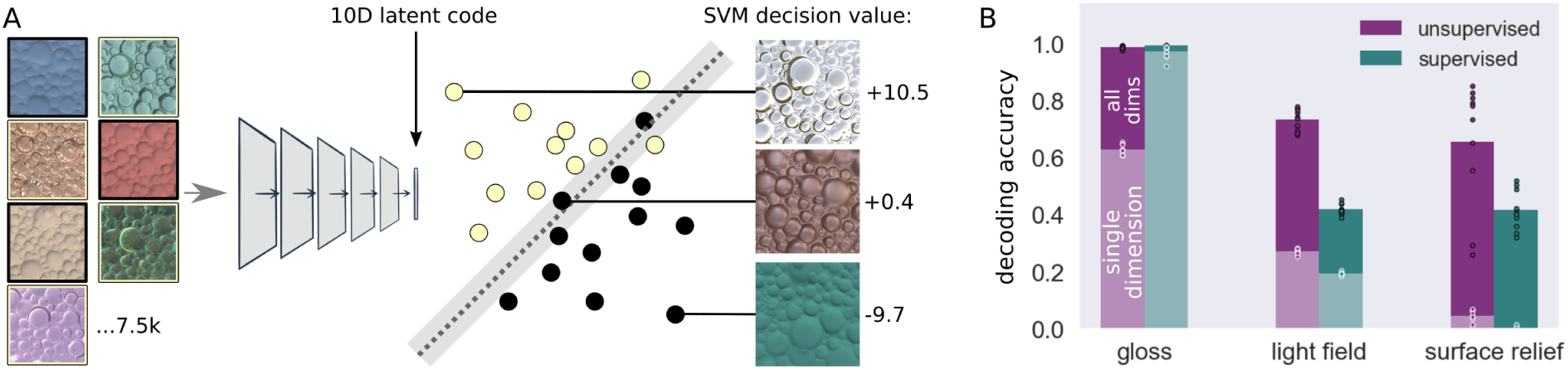
Disentangled world factors are available via a simple linear readout. (A) To quantify how explicitly gloss was represented in each model, we trained linear SVM discriminants to classify gloss from the representations of images within the latent code of each network. As well as obtaining decoding accuracy, this allowed us to use decision values (i.e. distance from SVM hyperplane) to make continuously-valued predictions from the model regarding the gloss of novel images (three examples shown), which was critical in subsequent experiments. (B) Average decoding accuracy for each world factor from latent codes of unsupervised and supervised networks (y-axis: proportion correct for two-way gloss classification and six-way light field classification, and *R*^2^ for surface relief regression). Pale lower regions of bars indicate average decoding accuracy when using the best single latent dimension, and give an impression of how distributed the representations are. Datapoints show individual performance for each of the ten training instances of each model type (black outline: full dimensionality, white outline: best single dimension).

Based on this linear decoding, we find that gloss-classification accuracy for novel renderings across the ten unsupervised models was extremely good at 99.3% (+/-SD 0.002)—practically as good as decoding gloss from the 10D latent code of the supervised models (99.4% +/- 0.002; Figure 3B). Light field and surface relief could also be decoded well above chance from the unsupervised networks (Figure 3B), and significantly better than from the supervised networks (independent-measures t-test comparing light field decoding between unsupervised and supervised models: *t_18_* = 23.25, *p* < 0.001, Cohen’s d = 10.40, 95% CI of difference = [0.28–0.34]; surface relief: *t_18_* = 3.30, *p* = 0.004, d = 1.48, 95% CI = [0.08– 0.36]). Thus, linear decoding further demonstrates that the unsupervised networks learn a compact representation that summarises information about not only surface material, but other scene properties such as illumination and surface relief. The analysis also revealed that representations were distributed rather than sparse. The full latent code predicted scene properties much better than any individual dimension could (Figure 3B and Supplementary Figure 1C).

Crucially, we could now derive a predicted gloss value for any image by inputting it to a network and calculating the SVM decision value for its corresponding point in latent space (i.e., signed distance of point from network’s gloss-separating hyperplane; Figure 3A). This allowed us to compare the model against human gloss perception.

For *<u>Experiment 1: Gloss ratings</u>* we rendered 50 new images of surfaces with random magnitudes of specular reflectance, sampled uniformly from almost matte to almost mirror-like. Twenty observers rated the apparent gloss of each surface, from 1 (matte) to 6 (glossy). We compared their ratings to the gloss values predicted by the unsupervised model. Figure 4A shows that agreement was excellent (mean *R*^2^ over ten model training instances = 0.84), and was substantially better than for the supervised model (mean *R*^2^ = 0.40; independent-samples t-test *t*_18_ = 12.45, *p* < 0.001, Cohen’s d = 5.57, 95% CI of difference = [0.37–0.50]). Notably, the unsupervised model even predicted human ratings better than ground truth (specular magnitude within the rendering engine; *R*^2^ = 0.73, one-sample t-test of difference, across model training instances *t*_9_ = 4.74, p = 0.001, d = 1.50, 95% CI = [0.05–0.13]).

**Figure 4:**
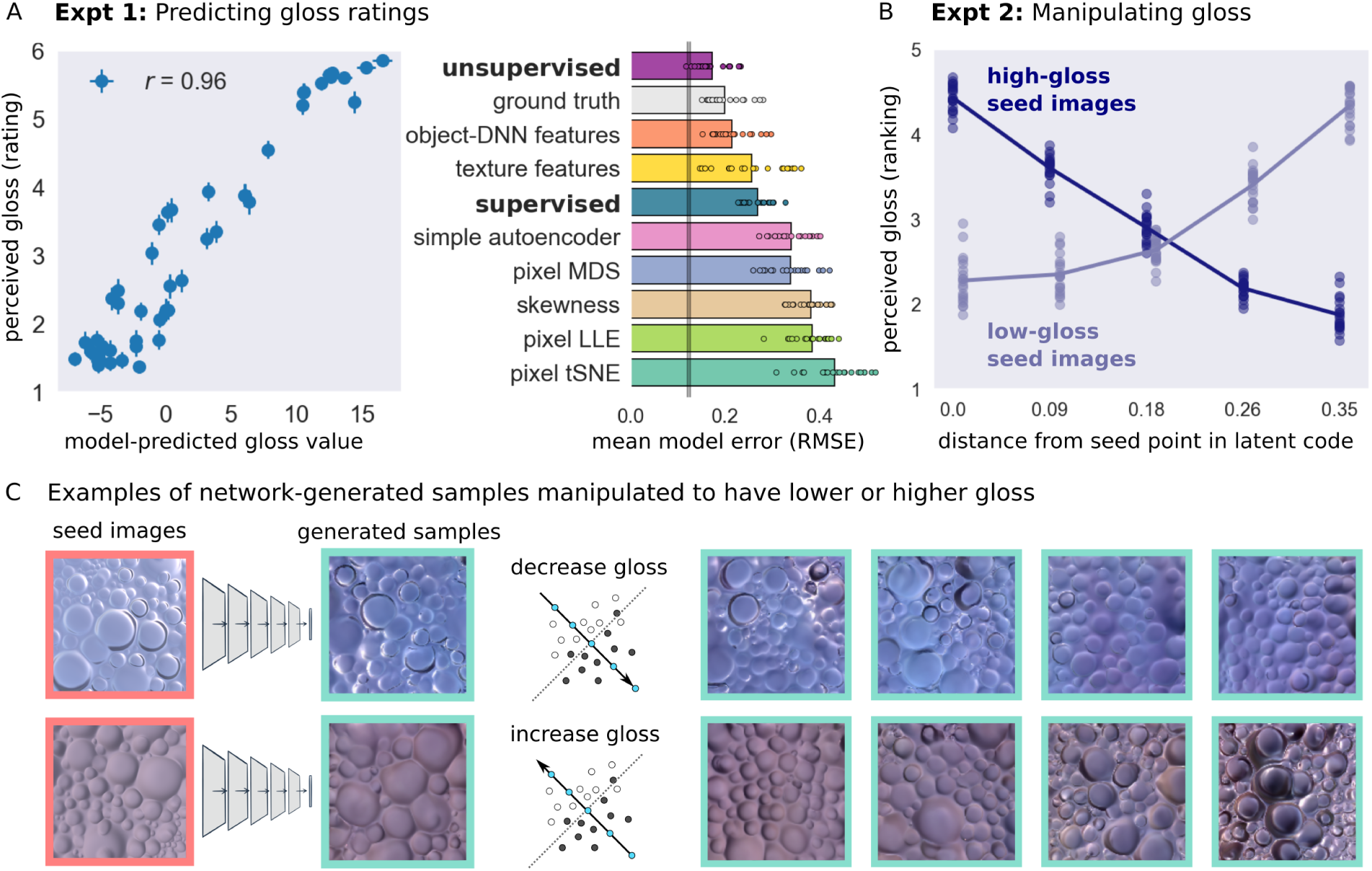
The unsupervised model can be used to modulate the gloss of generated samples, and predicts human-perceived gloss of new surfaces. (A) Gloss values in the unsupervised networks well predict human gloss ratings for 50 novel rendered surfaces of random gloss levels. Scatterplot shows decision value on a glossy-vs-matte SVM classifier for each image in the PixelVAE networks on the x-axis (averaged over ten training instances), and human gloss ratings on the y-axis (N=20). Error bars: SEM over 20 observers (vertical) or 10 model training instances (horizontal). Barplot shows average error in predicting each individual observer’s ratings, after normalising ratings and model predictions into the range 0-1, for the unsupervised model and for diverse alternative predictors (see text and Methods for details). Vertical grey line indicates how well individual human ratings can be predicted from data of other observers, giving the minimum possible model error. Datapoints show values for each observer. (B) Moving along the gloss-discriminating axis in a network’s latent code as shown in C successfully modulates perceived gloss, both when adding gloss to initially matte surfaces (pale blue line; *F_4,76_* = 649.82, *p* < 0.001, η^2^ = 0.97, 95% CI *r* = [0.97-0.98]), and when removing it from initially glossy surfaces (dark blue line; *F_4,76_* = 244.11, *p* < 0.001, η^2^ = 0.93, CI = [0.87-0.93]). Dots show each observer’s gloss rankings of each of five modulation steps, averaged over four test sequences starting from low-gloss initial seed images, and four starting from high-gloss seed images. Lines indicate mean over 20 observers. (C) Example gloss-modulated sequences. A glossy or matte rendered image (left) is input to a network and its 10D latent representation recorded. New images are generated from the model, conditioned first on the original (“seed”) point in latent space, and after stepping by small amounts along the model’s gloss-discriminating axis (i.e. the axis orthogonal to the decision plane of a glossy-vs-matte SVM classifier), either towards the “matte” (top) or “glossy” (bottom) directions.

We also considered a number of alternative models (Figure 4A bargraph), all of which predicted human judgements less well than ground truth. The best of these was a feature space consisting of the 1,000 final-layer features from a ResNet DNN trained on 1.2 million images to classify objects ^58, 71^. This is consistent with previous findings that representations in object-recognition DNNs capture perceptually-relevant features of textures and images ^61, 82–84^. Other, less well-performing, models included a multi-scale texture description comprising 1,350 feature dimensions ^85^; the 4,096 latent features from a relatively simple image autoencoder; 10-dimensional embeddings of raw images via tSNE ^74^, MDS or LLE ^75^; and luminance histogram skewness—a measure previously proposed to predict human gloss perception ^86^ (see Methods). Supplementary Figure 2 shows visualisations of how gloss and other scene factors are organised within each of these feature spaces.

Since PixelVAE networks are generative models, novel images can be generated by sampling from them (Figure 1D). In *<u>Experiment 2: Gloss manipulation</u>*, we used such images to test whether perceived gloss varied systematically with an image’s location in the model’s latent space. From each of the model training instances, four sequences of five images were generated by conditioning the image-sampling process first on a ‘low gloss’ point in the latent space and then on progressively higher-gloss points (i.e., moving along the model’s ‘gloss discriminating axis’, orthogonal to the SVM hyperplane). Another four sequences progressed from high to lower gloss points (Figure 4C). The same 20 observers sorted the images within each sequence from least to most glossy. Figure 4B shows that moving along the model’s gloss discriminating axis systematically reduced the apparent gloss of generated images when moving in the matte direction (one-way repeated-measures ANOVA *F_4,76_* = 244.11, *p* < 0.001, η^2^ = 0.93, 95% CI of correlation *r* = [0.87–0.93]) and increased it when moving in the high-gloss direction (*F_4,76_* = 649.82, *p* < 0.001, η^2^ = 0.97, 95% CI *r* = [0.97–0.98]).

Thus, despite never being given information about scene properties during training, we find the unsupervised networks develop internal representations that not only disentangle distal causes that are impossible to tease apart in the raw input (Figure 2B and Supplementary Figure 2), but also, more remarkably, predict human gloss perception better than the true physical reflectance properties of surfaces.

### The unsupervised model predicts failures of human gloss constancy

Although human gloss perception generally aligns well with specular reflectance, it also exhibits some well-documented ‘errors’. For example, bumpier surfaces tend to look glossier than flatter surfaces, and specific combinations of lighting and surface relief yield specific patterns of misperception (Figure 5A; ^34, 35^). Mimicking such perceptual errors is a key test of any computational model of biological vision ^87^. We assessed how well different models capture the systematic failures of gloss constancy exhibited by human observers.

**Figure 5:**
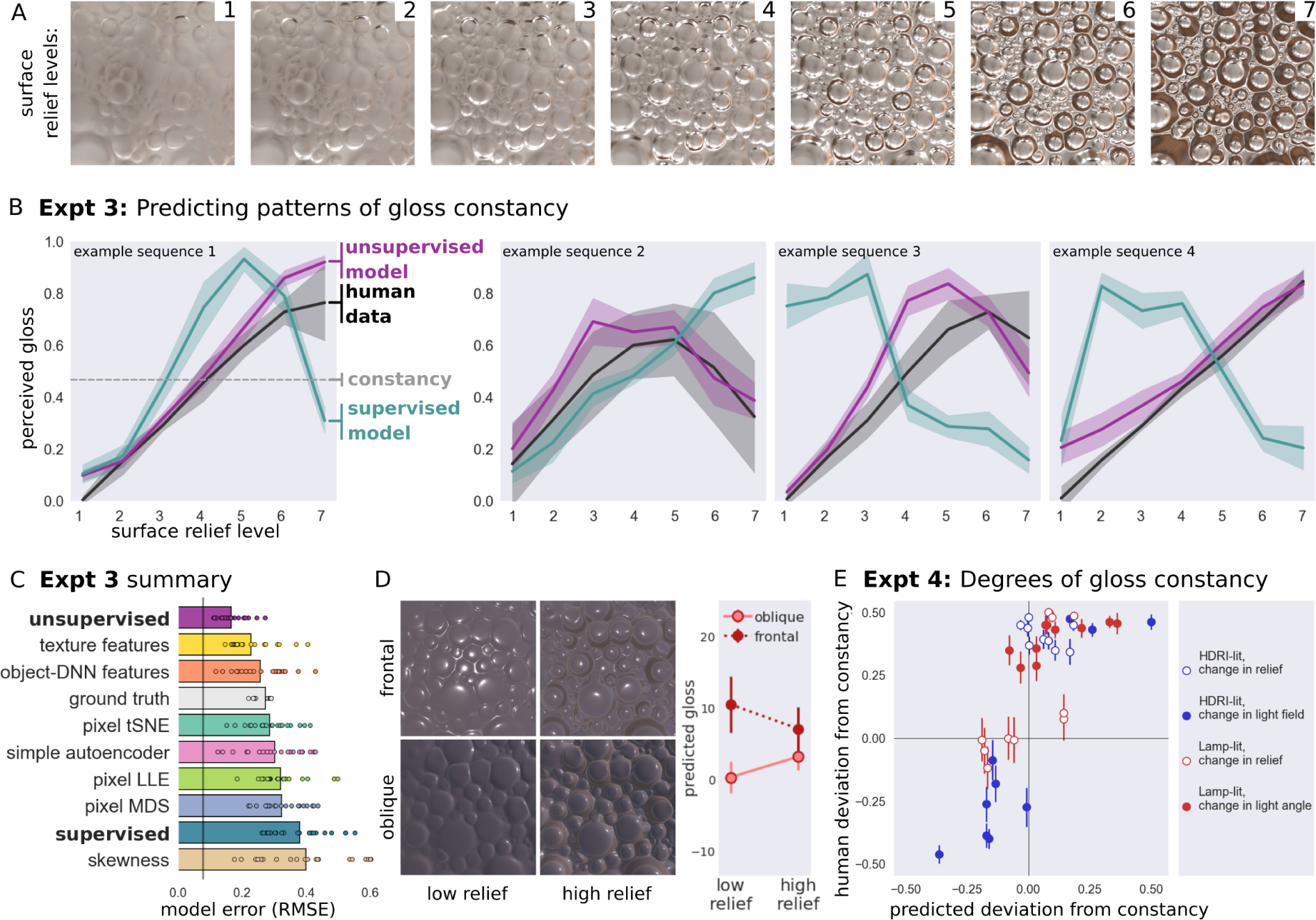
Unsupervised learning predicts human perception and misperception of gloss on an image-by-image basis. (A) Material properties of surface are identical in all seven images, but surface relief increases. People often see bumpier surfaces as glossier than flatter ones, constituting a failure of gloss constancy^34, 35^. (B) Examples of various patterns of human gloss constancy failure for four sequences of images increasing in surface relief. Leftmost panel shows data for sequence in (A). Black: human data (proportion of times each image step was reported as being glossier than others within each sequence); Purple: prediction of unsupervised model; Teal: prediction of supervised model. Veridical gloss perception would be flat horizontal line. Shading indicates standard deviation over 20 human observers or 10 model instances. See Supplementary Figure 3 for full data. (C) Summary of average error of each model’s prediction of each observer’s data, averaged across all twenty image sequences. Conventions are as in Figure 4B. (D) Changes in lighting also often change apparent material properties for human observers. The quartet of images shows the same surface material rendered on a low- vs high-relief surface (left vs right columns), and illuminated straight-on vs at a 30-degree angle (top vs bottom rows). Right: unsupervised model predictions for these four images (error bars: standard deviations over ten training instances). The model predicts a larger difference in apparent gloss for the low-than high-relief images, and an interaction between changes in lighting and relief. (E) Magnitude of deviations from constancy for 40 pairs of images depicting the same surface material with different scene properties (proportion of times image A of a pair was seen as glossier than image B). Scenes were illuminated by either natural lighting (blue dots) or directional lamp (red dots). Image pairs differed in surface relief (unfilled dots), light field (blue filled dots), or lighting angle (red filled dots). Y-axis: average human data over 20 observers, expressed in terms of raw proportion minus 0.5, so that zero indicates good constancy; +0.5 indicates image A always ranked glossier than image B; −0.5 indicates opposite. Error bars: SEM over observers. X-axis: difference in predicted gloss values between the images in each pair, averaged over 10 training instances of the PixelVAE model.

To do this, we rendered sequences where surface relief increased in seven steps, while reflectance and other scene properties remained fixed (Figure 5A). For stimuli in *<u>Experiments 3a and 3b: Patterns of gloss constancy</u>* we selected two sets of 10 sequences for which (a) both the unsupervised and supervised models predicted deviations from constant gloss, and (b) the models made *different* predictions about the particular pattern of deviations (see Methods). The rationale behind this is that cases where models disagree provide the strongest power to test which model is superior ^88, 89^.

With these image sequences in hand, in *<u>Experiments 3a and 3b: Patterns of gloss constancy</u>*, two groups of 20 observers judged gloss in a paired-comparison task (see Methods). If observers correctly estimated reflectance, all surfaces should appear equally glossy, yet we find that they do not. Observers exhibited strong failures of gloss constancy, usually reporting surfaces with deeper relief to be glossier, although perceived gloss was non-monotonic in seven of the twenty sequences, being highest for intermediate reliefs (four examples shown in Figure 5B; complete data: Supplementary Figure 3). The unsupervised model, despite never being explicitly trained to represent gloss, and without being fit to human data, predicted patterns of failures of gloss constancy remarkably well (median *R*^2^ across sequences and model training instances = 0.71). The model correctly predicted the qualitative pattern of constancy failure (monotonic vs non-monotonic) for 18 out of the 20 stimulus sequences. In contrast, the supervised model completely failed to predict human gloss constancy. For almost all sequences it made predictions that were anti-correlated with human patterns (median *R*^2^ = −1.45). Of the alternative models (Figure 5C), mid-level texture features ^85^ provided the next best performance (median *R*^2^ = 0.54), but was significantly poorer than the unsupervised model (one-sample t-test of difference across model training instances *t_9_* = 10.48, *p* < 0.001, Cohen’s d = 3.31, 95% CI of difference = [0.14–0.20]).

Human gloss constancy ranges from good to bad depending on interactions between lighting and shape ^30, 33–35, 90^. For *<u>Experiment 4: Degrees of gloss constancy</u>*, we rendered 40 image pairs depicting surfaces with identical material but different surface relief or lighting (Figure 5D; Methods), for which the unsupervised model predicted a wide range of degrees of gloss constancy, from excellent (near-identical predicted gloss for both images within a pair) to very poor (images received very different predicted gloss values; see Methods).

Twenty observers indicated which surface in each pair appeared glossier, with each pair repeated eight times. The unsupervised model predicted the degree and direction of human (failures of) constancy reasonably well (mean *r* across model training instances = 0.70, Figure 5E) and outperformed all alternative models (next-best model object-DNN features *r* = 0.64; one-sample t-test of difference, across PixelVAE training instances *t_9_* = 4.00, *p* = 0.003, Cohen’s d = 1.27, 95% CI of difference = [0.03– 0.08]). Fitting a simple logistic function to relate model and human gloss values further improves the prediction (from average *R*^2^ across model training instances = 0.60 for a linear fit, to *R*^2^ = 0.74 for a logistic fit).

Aggregating results across the three experiments using renderings (Experiments 1, 3, and 4), the unsupervised model predicts human perceptual judgements better than all others (Figure 6). It achieves near-perfect ground-truth gloss-classification accuracy, while still predicting idiosyncratic errors of human gloss perception (Figure 6A), with a feature space two orders of magnitude more compact than the next best model (Figure 6B). The next best model was a set of texture features hand-engineered to efficiently capture higher-order statistical structure in images ^85^, which performed significantly less well (one-sample t-test of difference in RMSE predicting individual data across three experiments, *t_59_* = 10.73, *p* < 0.001, Cohen’s d = 1.39, 95% CI = [0.05–0.07]). Simple image statistics, such as the skewness of the luminance histogram ^86^ failed to capture human gloss perception under our deliberately challenging stimulus conditions, where differences in specular reflectance must be de-confounded from differences in lighting and surface shape.

**Figure 6:**
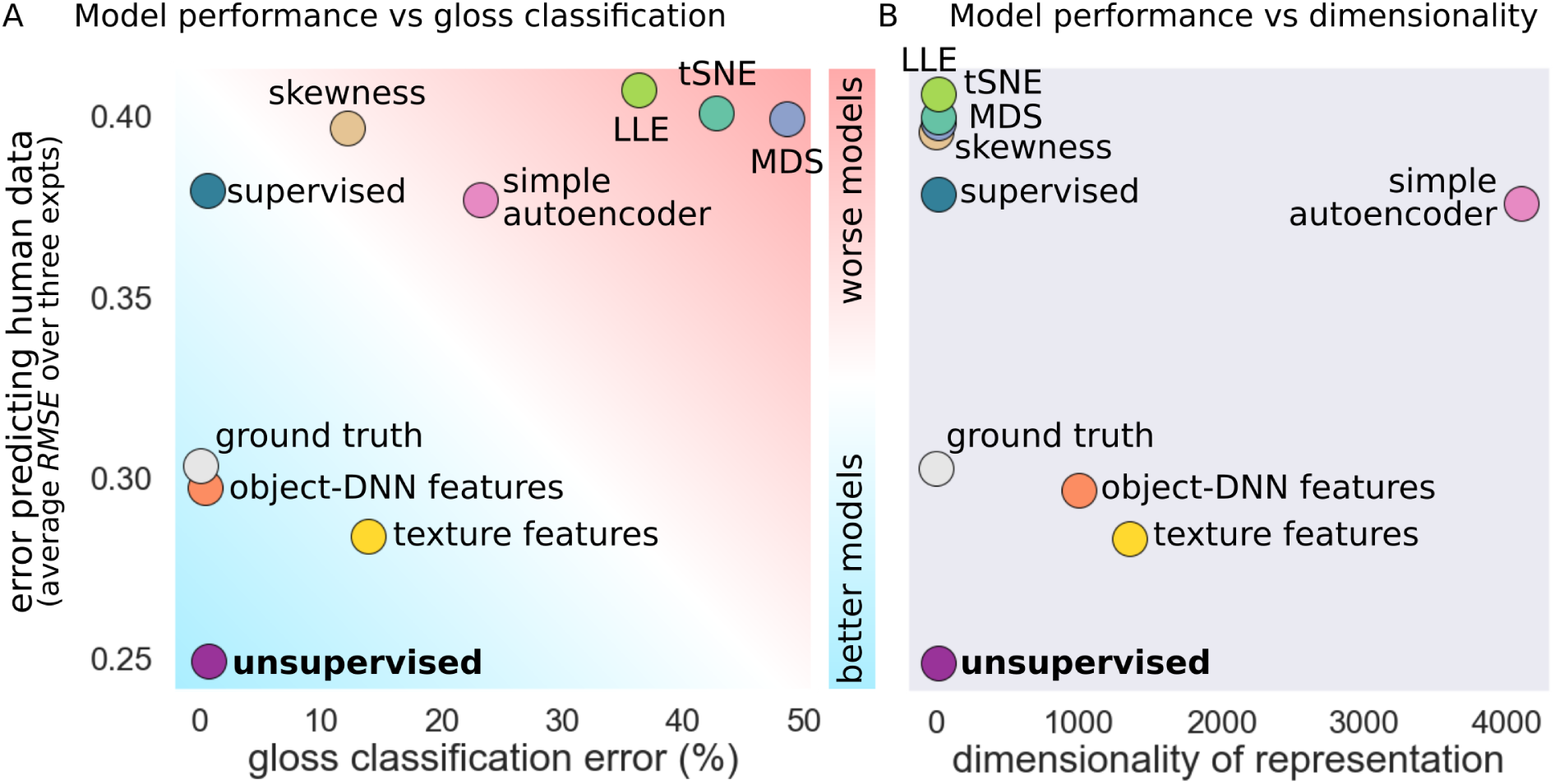
The unsupervised model outperforms diverse alternative models. (A) Relationship between error at classifying surfaces as glossy or matte (x-axis) and error in predicting human psychophysical gloss judgements (y-axis) for all models. Data are averaged over the three psychophysical experiments in which the ground truth gloss of stimuli is defined (Experiments 1, 3 and 4, each with N=20). (B) Relationship between the dimensionality of each model’s representation, and its error in predicting human judgements (y-axis shared with A). Not only does the unsupervised PixelVAE model best explain human data (difference from next-best model: *t_59_* = 10.73, *p* < 0.001, Cohen’s d = 1.39, 95% CI = [0.05-0.07]), but it does so with around two orders of magnitude fewer dimensions than the nearest competing models (a 1,350-dimensional texture feature space^85^, and a 1,000-dimensional layer from an object-recognition supervised DNN^57^).

### Model generalization and effects of training set

The composition of the training dataset has profound effects on all machine learning models. However, a good model of human mid-level vision should generalise over changes to training or test data. Our unsupervised networks were trained on a simulated environment with bimodally distributed gloss (near-matte or high-gloss). Nevertheless, they well predicted gloss in new image sets containing continuously-varying gloss (Figure 7A, mean *R*^2^ = 0.79 +/- SD 0.03; see also Supplementary Figure 6A), and predicted gloss for scenes with novel geometries and light fields as well as they did for familiar scenes (Figure 7A).

**Figure 7:**
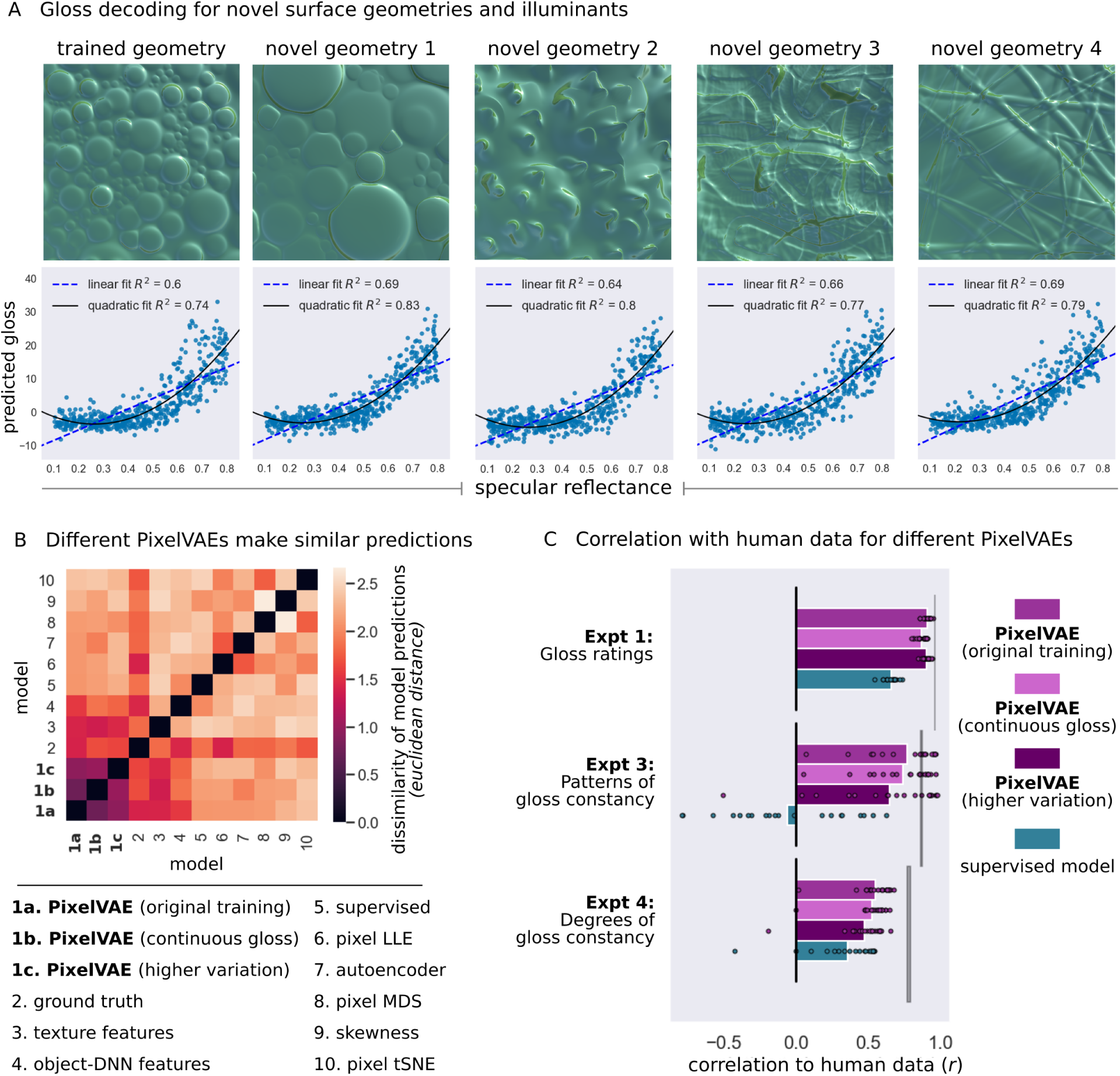
The unsupervised model generalises well to novel scenes, and makes similar gloss predictions after retraining on different visual environments. (A) Scatterplots show predicted gloss (y-axis) and ground-truth specular reflectance (x-axis) for sets of 500 new images rendered with random continuously-sampled gloss levels, using the trained surface geometry (leftmost panel) or four novel geometries unseen during training (right panels). A single example image from each set is shown above plots (magnitude and concentration of specular component = 0.65). All scenes were illuminated by light fields never seen during training. (B) New instances of the PixelVAE model with substantially different visual experience during training all make similar predictions for the gloss levels of experimental stimuli. Matrix shows the dissimilarity between the gloss predictions made by (1a) the original set of ten PixelVAE networks, averaged over training instances; (1b) five PixelVAE networks trained on a new set of 10,000 images using the same surface geometry and light fields, but with specular reflectance continuously sampled, averaged over training instances; (1c) a PixelVAE network trained on a third set of 10,000 images using ten surface geometries and 50 light fields, non-overlapping with the geometry and light fields used in the main training dataset; and (2-10) alternative models, ordered by the similarity of their predictions to that of the original PixelVAE. Gloss predictions are for all stimuli used in Experiments 1, 3 and 4, and were normalised into the range 0-1 for comparability across models. (C) Average correlation between model-predicted gloss and individual human judgements in each psychophysical experiment for PixelVAE implementations described in B. Datapoints show values for each observer (N=20 in each experiment). All unsupervised models predicted human data better than the supervised model in all experiments (across nine comparisons: *t*_19_ = 4.01–31.08, *p* < 0.001, Cohen’s d = 0.56-7.00, 95% CI = [0.06-0.18]-[0.60-1.07]). Vertical grey lines indicate how well data can be predicted on average from that of other observers, giving the highest possible average model correlation.

Model predictions do not seem to depend on artefacts of the specific computer graphics techniques used to generate our images (realtime rasterised rendering), as gloss predictions were near-identical for matched surfaces rendered using more time-intensive but physically faithful raytraced rendering (Supplementary Figure 4B). Remarkably, given their constrained training environments, the models even seem able to broadly categorise close-up photographs of real-world surfaces ^91^ as being of high or low gloss, although fail when shown surfaces far outside their training data, such as fabrics with high-contrast patterns (Supplementary Figure 6C-D).

We performed two tests of robustness to different training datasets. First, five new PixelVAE networks were trained on 10,000 additional renderings in which gloss was sampled continuously rather than bimodally (“continuously-sampled gloss training dataset”; Methods). Second, an additional PixelVAE was trained on a third dataset of 10,000 renderings in which surface geometry and lighting varied far more widely (“higher-variation training dataset”; Methods), with each scene comprising one of 10 novel surface geometries combined with one of 50 novel light fields. Both new training environments produced models with latent codes that could well subserve gloss classification on the original bimodal gloss dataset (mean accuracy = 96.7% and 91.4%, respectively). Importantly, we found that all three versions of the PixelVAE model (original, continuous-gloss training, and higher-variation training) made highly similar predictions regarding the relative gloss levels of experimental stimuli. This indicates that the ability to predict human perception is not highly sensitive to training set. The three versions of the unsupervised PixelVAE model, trained on non-overlapping datasets, made more similar gloss predictions to one another than to those made by any of the ten diverse alternative models (dark cluster of low dissimilarity values in the bottom left of Figure 7B). All three training environments led to unsupervised models that predicted human data reasonably well (Figure 7C), both for gloss ratings of novel rendered images (Experiment 1), and for the more challenging task of predicting patterns (Experiment 3) and degrees (Experiment 4) of (failures of) gloss constancy. Each of the three model versions predicted human data significantly better than the supervised model in all experiments (*t*_19_ = 4.01–31.08, *p* < 0.001, Cohen’s d = 0.56–7.00, 95% CI = [0.06–0.18]–[0.60–1.07] in nine repeated-measures t-tests comparing unsupervised vs supervised model correlation with individual participants’ data; bonferroni-corrected alpha = 0.006), and were significantly better than the most promising alternative model, texture features (80), in six out of nine comparisons (*t*_19_ = 0.73–5.84, *p* = 0.47–<0.001, d = 0.17–1.16, 95% CI = [-0.12–0.25]–[0.12–0.31]).

Several analyses were performed to assess how robustly unsupervised models outperformed their supervised counterparts. In building and training a DNN, values must be chosen for the many hyperparameters controlling network architecture and training. We evaluated the effects of some of these hyperparameters by training 28 additional models (14 unsupervised and 14 supervised), that differed from the original implementations in depth, learning rate, learning rate decay, training batch size, and complexity of the learned model (for unsupervised PixelVAEs); see Supplementary Table 1 for details. Eleven of the 14 unsupervised network variants outperformed all supervised network variants in predicting human gloss judgements; the only exceptions were networks that failed to train due to poor learning rate settings (Supplementary Figure 5). We also found that representations in the unsupervised model better predicted human judgements than those in the supervised model for all intermediate layers (Supplementary Figure 4A). Finally, we created a version of the supervised model that outputs continuous-valued gloss estimates rather than categorical decisions and trained it on a dataset with continuous rather than bimodal reflectances (see Supplementary Methods and Results). This version better predicted human gloss judgements than the category-supervised model, but less well than the unsupervised model, failing to exhibit the systematic *errors* that characterize human gloss perception (Supplementary Figure 4B). Overall, unsupervised learning in PixelVAE models appears to converge on a representation that captures key aspects of human gloss perception, and tolerates changes in the particular network hyperparameters, or the statistics, illuminations, or geometries of the training and test sets.

### Features underlying gloss representation in the model

Previous research ^35^ identified specific image features—the coverage, contrast, and sharpness of specular highlights—that predicted perceived gloss for surfaces like those evaluated here. To test whether the PixelVAE model was also sensitive to these cues, we measured the coverage, contrast, and sharpness of highlights in 10,000 new renderings of surfaces with specular reflectance varying continuously from near-matte to near-mirror. All three cues could be decoded from the latent code of a PixelVAE trained on these images (mean *R*^2^ = 0.71), and could be increasingly well decoded from successive convolutional and fully connected layers (Supplementary Figure 7A). A linear combination of the three cues in the layer immediately preceding the 10D latent code correlated with gloss predicted from the latent code (*r* = 0.80; Supplementary Figure 7B). Moreover, manipulating the highlights in images to weaken each cue also reduced predicted gloss (Figure 8A-B; one-way repeated-measures ANOVA for the effect of highlight contrast reduction: *F_9,81_* = 11.20, *p* < 0.001, η^2^ = 0.55, 95% CI of correlation *r* = [-0.65–-0.41]; sharpness: *F_9,81_* = 9.65, *p* < 0.001, η^2^ = 0.52, 95% CI *r* = [-0.61–-0.40]; coverage: *F_9,81_* = 18.14, *p* < 0.001, η^2^ = 0.67, 95% CI *r* = [-0.58–-0.29]).

**Figure 8:**
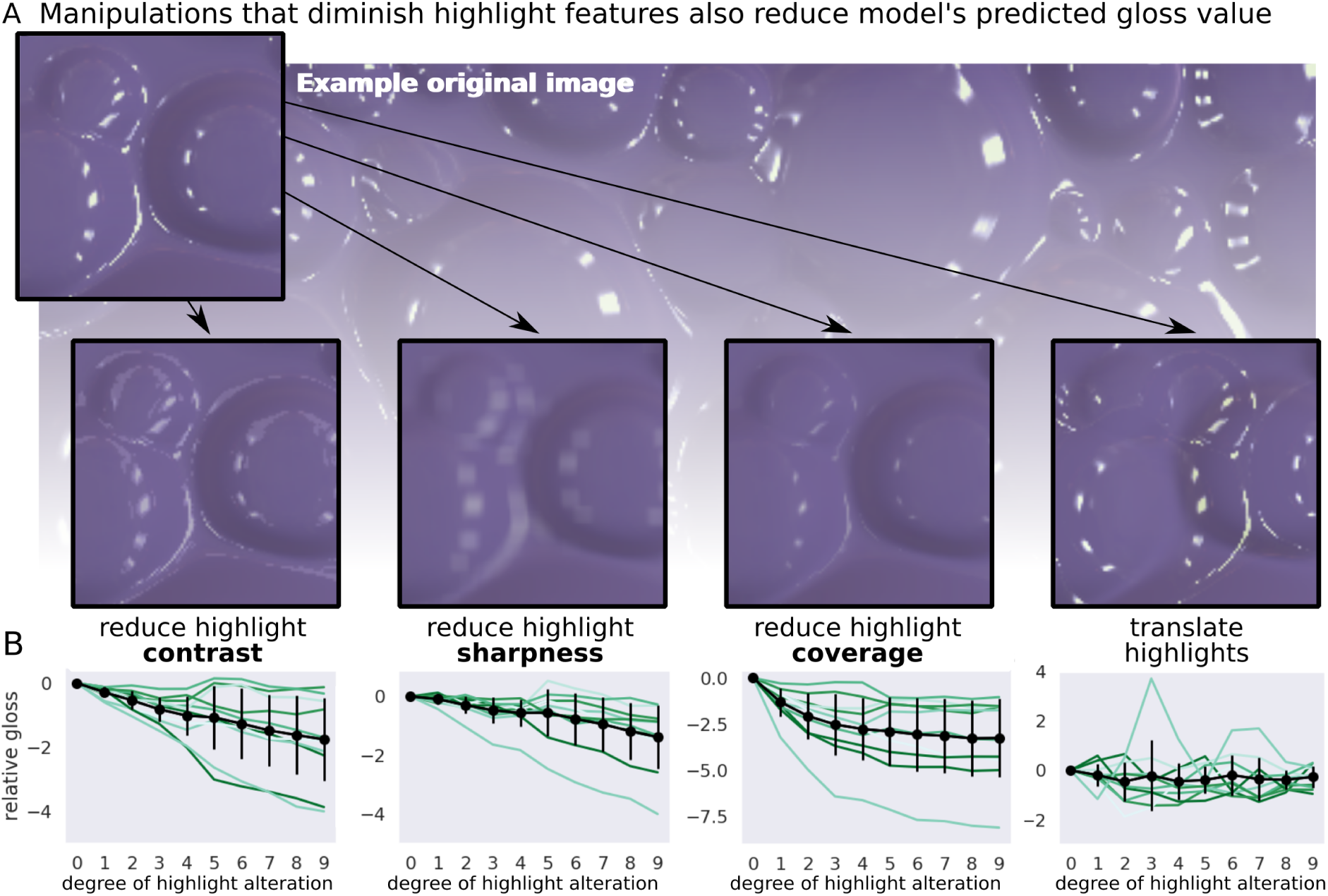
Reducing the coverage, sharpness or contrast of highlights reduces model-predicted gloss. (A) We separately rendered the specular and matte components for ten probe images, and manipulated the specular component by either reducing contrast, blurring, shrinking coverage via an image erosion algorithm, or laterally translating it, in ten progressive steps. We then recombined the two components to produce ten sequences of images, and presented them to the unsupervised networks. Insets show each of these manipulations applied to part of an example image. (B) Model-predicted gloss declined when any of the three highlight features was reduced (left three plots; contrast: *F_9,81_* = 11.20, *p* < 0.001, η^2^ = 0.55, 95% CI *r* = [-0.65–-0.41]; sharpness: *F_9,81_* = 9.65, *p* < 0.001, η^2^ = 0.52, CI = [-0.61–-0.40]; coverage: *F_9,81_* = 18.14, *p* < 0.001, η^2^ = 0.67, CI = [-0.58–-0.29]). Green lines show the predicted gloss for each step in each sequence, relative to that of the original un-manipulated image, averaged over ten training instances of the model; black lines show mean and standard deviation over all sequences. Translating the specular components (rightmost plot), which does not affect the coverage, sharpness, or contrast of highlights, did not measurably affect predicted gloss (*F_9,81_* = 0.48, *p* =0.88, η^2^ = 0.05, CI = [-0.21–0.09]).

Interestingly, we did not find evidence that the model’s predicted gloss decreased when the specular component of images was shifted so that highlights were misaligned with the geometry of the depicted surfaces (Figure 8B; *F_9,81_* = 0.48, *p* =0.88, η^2^ = 0.05, 95% CI *r* = [-0.21–0.09]). This manipulation preserves coverage, contrast and sharpness, yet infringes photogeometric constraints that are a precondition for humans to identify highlights as specularities, and therefore to see gloss at all ^6, 92, 93^. The fact that the model predicts human gloss constancy without appearing to be sensitive to such constraints suggests that although these constraints are crucial to many aspects of surface perception ^7, 93–96^, the degree of perceived gloss, within surfaces that are seen as having highlights, may be largely explainable in terms of image features.

## Discussion

Efficient representation of sensory data has long been hypothesised as a central goal of neural coding ^15, 19–24, 97^. Yet, while such approaches predict many aspects of low-level image encoding, to date, they have not explained how we visually infer properties of the outside world. Unsupervised learning objectives in modern DNNs, such as data compression and spatial prediction, offer powerful new implementations of these statistical learning principles ^17^. Our findings show that mid-level perceptual dimensions, like gloss—which imperfectly map onto properties of the physical world—can emerge spontaneously by learning to efficiently encode images. Thus, unsupervised learning may provide a bridge that links theoretical views emphasizing the importance of image statistics (e.g. ^86, 98, 99^), to those that treat perceptual processes as a decomposition of images into distinct physical causes (e.g. ^4, 6, 8, 100^).

One of the fundamental unsolved questions in vision science is how the visual system became aware of the different physical sources that contribute to image structure. Perception is commonly framed as the optimal *estimation* of a set of known physical quantities ^4, 9–11^. But these quantities that the brain putatively estimates were not specified *a priori*; they must somehow be discoverable (over either the course of evolution or learning) based on their manifestation in sensory experience ^12–14^. Here, we suggest that different physical causes give rise to different high-order regularities in visual data, making them discoverable through data-driven unsupervised learning processes ^15–18^. We provide a proof-of-principle that it is possible to learn to disentangle distal causes without prior knowledge about which classes of causes exist in the world, the cues that could be used to distinguish them, or even how many different classes of causes there are. An unsupervised statistical learning predicted both the expected changes in perceived gloss caused by varying specular reflectance (Experiment 1; ^32, 33, 39–41, 43^), as well as illusory changes in perceived gloss that arise from varying lighting and shape (Experiments 3 and 4; ^34, 35^). We suggest that known systematic errors in gloss perception can be attributed to the particular pattern of partial disentanglement arising from unsupervised statistical learning of surface appearances.

One of our more intriguing results is that the unsupervised model predicted human perception better than the supervised model tested. It is important to note that this is not because humans and unsupervised networks were better at extracting ground truth in these stimuli. Categorisation-supervised networks categorised gloss almost perfectly (Figure 3b), and regression-supervised networks predicted continuous gloss levels almost perfectly (Supplementary Figure 4b), yet both predicted human judgements less well than unsupervised networks. This implies that the systematic errors exhibited by humans and the PixelVAE model are not a trivial consequence of some inherent impossibility in recovering specular reflectance for these stimuli. Nor is it explained by the supervised model reporting ground truth specular reflectance *too* faithfully. Although the supervised model is near-perfect at coarsely categorising surfaces as having high or low specular reflectance, it still predicts different *degrees* of glossiness for different images, including sometimes erroneously. Yet the supervised and unsupervised models make *different* predictions on an image-by-image basis, with the latter more closely matching those made by humans. We propose that this shared pattern of deviation from ‘ideal performance’ may arise from shared characteristics in how the human visual system and unsupervised model learn to encode images.

One of the most notable failures of the PixelVAE in capturing human data is its insensitivity to photogeometric constraints known to affect human surface perception, such as the alignment of specular highlights with diffuse shading ^7, 92, 93^ (Figure 8). We believe that this failure is likely due to the relative poverty of 3D shape information in its training set. The link between highlights and diffuse shading arises from constraints imposed by the 3D shape of a surface ^94, 96^. It seems implausible to expect any visual system trained solely on monocular, static images to develop good sensitivity to these constraints, and without a detailed representation of 3D shape, no model is likely to explain all aspects of human gloss perception ^94, 95, 101^. We tailored the training sets towards modelling variations in perceived glossiness for physically realistic surfaces, where highlights are assumed to align with surface shading. ^32, 33, 39–41, 43^. An important direction for future research is testing whether unsupervised DNNs can also learn photogeometric relationships, if training sets provide additional information about shape (e.g., through motion, stereo, occlusion, or larger variations in geometry).

In using deep learning models, we do not wish to imply that all material perception is learned during an individual lifetime. Unsupervised learning principles can also operate on an evolutionary timescale. For example, V1 cell receptive fields are predicted by simple unsupervised learning models such as independent components analysis ^22^ and these seem to be present at birth in macaques ^102^. There is evidence that 5-8 month-old infants can distinguish between matte and specular surfaces ^103^, but also that material recognition and classification are still developing in 5-10 year-old children ^104, 105^. Even 5-8 months of visual experience provides a huge dataset for statistical regularity learning ^106^. It could be that approximate versions of many perceptual dimensions are rapidly learned during the first days, weeks and months of life.

Although we do not propose the PixelVAE architecture as a physiological simulation of visual cortex, its computational principles are eminently achievable in biological brains. Data compression and spatial prediction are learning objectives that require no additional information beyond the incoming sensory data, and there are several mechanisms by which brains could represent the probability distributions used within the PixelVAE network ^107–109^. At the same time, the brain certainly does not learn statistical distributions over RGB pixels. If the visual system uses learning objectives like the ones investigated here, they presumably operate on image representations that have undergone substantial retinal processing ^110, 111^.

In conclusion, unsupervised DNN models provide an ecologically feasible solution to the problem of how brains come to represent properties of the distal world without access to ground truth training data ^12–14, 17, 112^. Non-linear transformations reorganise inputs according to high-order regularities within and across images, allowing the visual system to better summarise and predict sensory data. Because regularities in images are caused by underlying physical objects and processes, these new configurations often end up (partially) disentangling physical properties from one another. Our results suggest that the imperfect nature of this disentanglement may account for the characteristic ‘errors’ humans make. Failures of constancy, which are rife in vision, may therefore offer tell-tale clues to how we learn to see. Unsupervised learning may account for them, not just in gloss perception but in perception more broadly.

## Methods

### Participants

Three groups of human naïve observers reported perceived gloss across five experiments: Experiments 1 and 2 (N=20, mean age 23.45 [range 19-32], 16 female, 4 male), Experiment 3a (N=20, mean age [range 19-31, 16 female, 4 male), Experiments 3b and 4 (N=20, mean age 24.45 [range 19-35], 14 female, 6 male). Six individuals participated in two different experimental groups, but received no information about the experimental design or hypotheses after the first session. No statistical methods were used to pre-determine sample sizes but our sample sizes are larger than those reported in previous publications measuring gloss constancy^34, 35, 37^. All participants had normal or corrected-to-normal visual acuity. Two male participants self-reported poor red-green colour vision. Experiments were conducted in accordance with the Declaration of Helsinki (6^th^ Revision), with prior approval from the ethics committee of Justus Liebig University, Giessen, Germany. Volunteers gave written informed consent and were paid €8 per hour.

### Stimuli

Stimuli were 800×800pixel images of bumpy surfaces rendered using Unity3D (version 2018.2.3). A virtual 40×40cm sheet with irregularly positioned bumps was placed in a scene and illuminated by one of six high dynamic range (HDR) light probes. Light probes were white-balanced 8192×4096pixel images of four exterior (beach, bay, woodland, savannah) and two interior (church, conference hall) environments, captured by Unity Technologies (https://assetstore.unity.com/packages/2d/textures-materials/sky/unity-hdri-pack-72511). A virtual camera (60° field of view) hovered 12cm above the sheet, viewing directly downwards. By randomly varying the camera’s location and orientation, an extremely large number of unique images could be generated from the scene.

#### Main training dataset

10,000 images were rendered to create the main training dataset for neural network models. For each rendering, one of the six HDRIs was randomly selected to illuminate the scene. Surface relief was scaled multiplicatively in depth by a uniformly-sampled random scaling factor, so that the distance between the lowest and highest points on the surface was between 0.1–2.5cm. The surface’s reflectance properties were controlled via the Unity3D standard shader, using the ‘specular setup’. The diffuse reflectance component of the surface material was chosen by selecting RGB values randomly uniformly within the interval 0.3 to 0.7, independently for each channel. The surface was either low or high gloss, with equal probability. For low gloss surfaces, the specular reflectance component of the material was selected randomly uniformly between 0.1–0.3, and the concentration of the specular component was randomly uniformly between 0.2–0.4. For high gloss surfaces, specular reflectance was between 0.3–0.5, and specular concentration between 0.75–0.95. The training dataset therefore had a bimodal distribution of approximately 50% low- and 50% high-gloss surfaces, with small variations in reflectance properties within each group. The same dataset was also used when training classifiers to decode gloss and other world factors from models.

#### Continuously-sampled gloss training dataset

To verify that the representations learned by our models were not artefacts of the bimodal gloss sampling in the main dataset, we rendered 10,000 new images in which both the magnitude and concentration of the specular reflectance component were sampled randomly uniformly between 0.1–0.8, independently of each other. All other scene properties were varied as in the main dataset.

#### Higher-variation training dataset

To verify that less constrained visual diets could lead to similar gloss representations, we created a third training dataset. 10,000 images each randomly combined one of ten novel surface geometries with fifty novel light fields. Novel geometries were virtual 40 ×40cm sheets with different shapes and sizes of irregularly placed ridges, bumps, and indentations (examples in Figure 7A). Novel light fields were fifty 4096×2048pixel HDR images (25 exterior, 25 interior) from the HDRI Haven database (http://www.hdrihaven.com). The distance of the virtual camera above the surface varied randomly uniformly between 8–12cm to introduce scale variation. All other scene properties varied as in the main dataset.

#### Renderings with continuously sampled gloss levels (<u>Experiment 1: Gloss ratings</u>)

Stimuli were 50 new rendered images with specular reflectance chosen randomly uniformly between 0.2–1.0, and specular concentration set to the same value. Other attributes were varied as in the main dataset. For all rendered images used as stimuli in psychophysical experiments (Experiments 1, 3 and 4), the same geometry and set of light fields were used as in the main training dataset, but images were novel renderings unseen by any model during training.

#### Gloss-modulated images generated from PixelVAE models (<u>Experiment 2: Gloss manipulation</u>)

For each of the PixelVAE networks, we first determined the axis in 10D latent space along which gloss could be most strongly discriminated (see **Data analysis**). Eight images (four low-gloss, four high-gloss) from the main training dataset were input to each network, and their corresponding latent points were recorded. These were used as ‘seed points’ to generate eight ‘gloss-modulated’ sequences for each network. The first step in each sequence was generated by conditioning the model’s pixel sampling process on the respective seed point in latent space, and sampling a 128×128pixel image (see **Neural network architectures and training**). The conditioning point was then moved 0.07 units in latent space along that model’s gloss-discriminating axis, either in the matte (for high-gloss seed images) or glossy (for high-gloss seed images) direction, and a second image was generated. This was repeated three more times, yielding five-step sequences for each seed image from each model training instance (400 experimental images in total).

#### Sequences of renderings with increasing surface relief (<u>Experiments 3a and 3b: Patterns of gloss constancy)</u>

To create a strong test of the different models, we wanted to probe human gloss constancy using stimuli (a) for which there were clear differences between constancy patterns predicted by unsupervised vs supervised models ^88, 89^, and (b) which were likely to produce diverse patterns of failures of constancy in human observers. We therefore first generated candidate stimuli, then selected those that best satisfied these desiderata.

For each experiment, we rendered 200 different candidate sequences of 7 images with fixed random material, illumination, and camera position, but exponentially increasing surface relief (maximum bump heights of {0.10, 0.18, 0.31, 0.55, 0.97, 0.70, 3.0}cm). All surfaces had relatively high gloss. In Experiment 3a, specular reflectance was randomly uniformly between 0.2–1.0, and specular concentration 0.7–1.0. In Experiment 3b, reflectance was between 0.1–0.3, and concentration 0.8–1.0. Other attributes were varied as in the main dataset. All 1400 images in each set were input to each of the unsupervised PixelVAE and supervised ResNet networks, to obtain predicted gloss values (see **Data analysis**). For each candidate sequence of 7 images, we then simulated the responses from each network under a two-alternative forced-choice (2AFC) experiment in which each possible pair of images was compared, and the image for which the model predicted a higher gloss value was selected. This was analogous to the “which is more glossy?” task performed by human observers (see **Psychophysical procedures**).

First, for each sequence we performed a 2×7 (model × surface relief) ANOVA between unsupervised and supervised model predictions, averaged over training instances of each model. Sequences were sorted according to the *F* interaction statistic, prioritising those with strong disagreements between model predictions ^88, 89^. In Experiment 3a, we visually inspected the top-ranked sequences and selected 10 with diverse appearances and subjective failures of gloss constancy.

In Experiment 3b we selected test sequences in an entirely automated manner by classifying sequences into four qualitative groups based on the average constancy pattern predicted by the unsupervised models:

- Group 1: model predicts approximate constancy (range of predicted “proportion glossier” values < 0.25, and a linear function fit achieved *R*^2^ > 0.70);
- Group 2: model predicts an approximately linear increase in gloss with bump height (range > 0.50, and a linear fit achieved *R*^2^ > 0.90);
- Group 3: model predicts a non-monotonic failure of constancy (range > 0.50, and a quadratic fit of *R*^2^ > 0.90 with a negative coefficient of the squared term);
- Group 4: model predicts an upward-swinging non-linear failure of constancy (range > 0.50, and a quadratic fit of *R*^2^ > 0.90 with a positive coefficient of the squared term).

We then selected the top sequences in each group (ranked by *F* interaction term), in proportion to the size of each group (1 “constant,” 4 “linear,” 2 “non-monotonic,” and 3 “non-linear” sequences).

#### Pairs of renderings with differing surface relief or lighting (<u>Experiment 4: Degrees of gloss constancy</u>)

Here we sought pairs of stimuli (a) with diverse scene differences (some differing in surface relief, some in lighting environment, and some in angle of illumination), and (b) for which the unsupervised model made a wide range of predictions, ranging from good constancy to strong failure of constancy.

We rendered 800 candidate image pairs with the same random material properties and camera position, but differing in either surface relief (0.40 vs 1.5cm maximum bump height) or illumination. Pairs differing in illumination were either ‘naturally’ lit (differing in HDR lighting environment, from among the six used in the ***main training dataset***), or illuminated by a directional lamp (at a 30° vs 90° angle). Candidate pairs were generated in four groups of 200:

- Group 1: identical natural light field; different surface relief
- Group 2: identical lamp angle; different surface relief
- Group 3: identical surface relief; different natural light field
- Group 4: identical surface relief; different lamp angle

Average predicted gloss values were obtained for each image, across training instances of the unsupervised PixelVAE model (see **Data analysis**). Within each group, we ranked pairs by the absolute difference in predicted gloss of the images in the pair, and selected pairs lying at each of the 10th percentiles, yielding 40 test pairs.

#### Geometry and lighting generalisation test set

To test generalisation of models trained on the main dataset, we rendered five sets of 500 new images, one using the original surface sheet, and four using novel surface geometries (Figure 7A). Illumination was randomly one of eight novel 4096×2048pixel HDR light probes (six exterior, two interior) from the Dutch Skies 360° HDRI Project (http://www.dutch360hdr.com/). Specular reflectance was varied as in the ***continuously-sampled gloss training dataset***.

#### Highlight-feature manipulated images

To manipulate highlight features (Figure 8), we manipulated the specular component image in four different ways, before combining with the diffuse component image by addition:

- (1) Translated highlights: specular component was shifted rightwards in ten 5-pixel steps. To avoid edge artefacts after translation, images were rendered at 1024×1024 resolution, and then cropped to the lower right 800×800 pixels.
- (2) Reducing highlight coverage: we applied image erosion to the specular component using the OpenCV package for Python. Kernel size was 2×2. Up to ten iterations were applied to create ten progressively reduced highlight maps.
- (3) Reducing highlight sharpness: specular image was convolved with a Gaussian filter, with step size ranging from 1×1 to 11×11.
- (4) Reducing highlight contrast: we created a ‘highlight mask’ by identifying pixels with values >4 (from 0-255 range) in greyscale version of specular image. Standard deviation of pixel intensity within the highlight map was multiplicatively reduced in ten steps, from 1 to 0.1 times their original SD, while retaining the original mean.

### Neural network architectures and training

DNNs were implemented and trained in Python 2.7 with Tensorflow 1.14 on a GPU-accelerated machine using 1-4 GTX1080 graphics cards. Networks were trained on the first 9,000 images of the ***main training dataset***, using the next 500 for validation, and the final 500 for testing accuracy of the supervised networks. All images were input at 128×128 pixel resolution. For both unsupervised and supervised models, ten independent instances of the same architecture were trained from different random initial weights and with different random sampling of training batches, to ensure robustness to representational differences between training instances ^70^. Additionally, we trained five instances of the PixelVAE network on the ***continuously-sampled gloss*** dataset and one on the ***higher-variation*** dataset (see **Stimuli**), dividing training and validation data in the same proportions. No data augmentation was used. All architectures used rectified linear activation functions.

#### Unsupervised PixelVAE model

We used the implementation from ^47^ of the PixelVAE architecture ^48^, available at https://github.com/ermongroup/Generalized-PixelVAE. The architecture consists of two streams of convolutional layers, which learn jointly via backpropagation. One stream is a ‘conditioning network,’ which is a convolutional variational autoencoder that takes an image as input and outputs a 10-dimensional smooth latent representation. We chose a 10D latent code as being approximately the most compact representation that still allowed the network to learn a good model of the training images, based on pilot data (see Supplementary Figure 1A-B). The other stream is an autoregressive PixelCNN++ ^113, 114^ model that learns the structure of the training data in terms of a polynomial probability distribution over pixel values, and takes as inputs both the image and the latent code output by the conditioning network. To generate new images, the autoregressive stream chooses an RGB value for each pixel, working from the top left to the bottom right of the image. Each pixel is sampled from the learned probability distribution, conditioning on both the values of pixels generated so far (which constrain the local structure of the image), and on the values provided in the latent code (which constrain more holistic image properties).

The conditioning network consisted of three convolutional layers of 64, 128, and 256 feature maps, followed by a fully connected layer of 512 units and a 10-unit latent code layer. The autoregressive network consisted of six ‘residual blocks’ of three layers of 64 convolutional feature maps each, with skip connections linking the first and sixth, and second and fifth blocks. Pixel likelihood distributions were modelled with a mixture of 12 logistic functions, and networks were trained with a batch size of 5 and a learning rate of 0.001 for 200 epochs, around which point validation error plateaued. Learning rate was gradually decayed by multiplying by 0.999995 after each epoch. No regularisation was used during training, to encourage the network to depend on information in its latent code ^47^.

#### Supervised ResNet model

We used the Tensorflow implementation of the ResNet architecture ^57^ from https://github.com/wenxinxu/resnet-in-tensorflow. Networks consisted of three ‘residual blocks’ each made up of three layers of 56 convolutional feature maps, with skip connections between each. The output of the final layer was passed to a 10-unit fully-connected layer, which we treated as the ‘latent code’ of the model for analysis purposes. This passed, via one more non-linearity to a two-unit softmax output layer. Ten networks were trained, from different random initial weights, to classify images as renderings of high or low specular reflectance surfaces (see **Stimuli**). Networks were trained with a batch size of 32 and a learning rate of 0.001 to minimise the sparse softmax cross-entropy between outputs and correct labels. Learning rate was gradually decayed by multiplying by 0.99995 after each epoch. Networks were trained for 21 epochs, by which point validation accuracy plateaued above 99%.

#### Simple autoencoder

We also considered a far simpler unsupervised model, in the form of a non-variational convolutional autoencoder implemented using Keras 2.2.5 for Tensorflow. The architecture comprised four layers of 64, 32, 16, and 16 feature maps alternating with 2 ×2 pooling layers, leading to a 4,096-unit fully connected bottleneck layer, which we treated as the model’s latent feature space for analysis purposes. The compressed code was expanded through mirrored layers of 16, 16, 32, and 64 feature maps to an output of the original dimensionality (128×128×3). The network was trained for 1,000 epochs to minimise mean absolute error between its input and output images (batch size 32, other learning parameters used default Adam optimiser values as implemented in Keras).

#### Object-trained deep neural network

Finally, we evaluated a pre-trained DNN supervised on object-recognition: an 18-layer Resnet ^57^ model available from the MATLAB 2019b Deep Learning toolbox (https://mathworks.com/help/deeplearning/ref/resnet18.html). The network had been pre-trained on the Imagenet Large-Scale Visual Recognition Challenge (ILSVRC) database to classify 1.2 million images into 1,000 object and animal categories ^58^. For analyses, we used representations in the 1,000-unit fully-connected layer immediately before softmax category readout.

### Additional comparison models

#### Histogram skewness

Skewness was defined as the skew (3rd moment) of the distribution of pixel intensities in a greyscale version of each image.

#### MDS / tSNE/ LLE

Multi-dimensional scaling (MDS) is a linear dimensionality reduction technique that finds the linear projection of a dataset that best preserves distances between all data points. t-distributed Stochastic Neighbourhood Embedding (tSNE ^74^) is a non-linear technique that preferentially preserves distances between nearby data points. Locally Linear Embedding (LLE ^75^) is a non-linear technique that seeks a lower-dimensional manifold that best preserves all distances.

Each dimensionality reduction algorithm was used to create a 10D embedding of the 10,000 images from the main training dataset, as well as all 270 rendered images used in experiments (50 images from Experiment 1; 140 from Experiment 3; 80 from Experiment 4). The experimental probe images were included because it is not possible to project new data points into the reduced-dimensionality solution discovered by tSNE or LLE. Additional 2D embeddings were performed to create visualisations (Figure 2A-C and Supplementary Figure 2) using 4,000 images randomly selected from the main training dataset. Default parameters were used, as implemented in the scikit-learn package for Python.

#### Texture features

For each image we calculated a multi-scale texture feature description ^85^ using the TextureSynth package for MATLAB (www.cns.nyu.edu/~lcv/texture/). Images were rescaled to 256×256 pixels, and analysed at 3 scales, 4 orientations, with a spatial neighbourhood of 7, producing a description of each image in terms of 1,350 feature dimensions.

### Psychophysical procedures

Psychophysical experiments were conducted in a dimly lit room, sitting at a comfortable distance from the screen. Participants could freely view the screen, and were given no fixation instructions. Stimuli were presented on an EIZO ColorEdge CG277 self-calibrating LCD monitor with a resolution of 2560×1440 and a refresh rate of 60Hz. PsychoPy 3.1 with Python 3.6 was used to present stimuli and record responses. At the beginning of each experiment, observers were shown 4-12 example experimental stimuli, randomly arranged, to familiarise them with the appearance and range of material properties they would be asked to judge. Response times were unconstrained, and stimuli were displayed until a response was recorded.

#### Experiment 1: Gloss ratings

Experiment 1 measured gloss ratings for novel rendered images. Fifty 800×800 pixel (18.6×18.6cm) images were presented singly in the centre of the screen. Each was repeated three times, for a total of 150 trials, presented in a random order. Participants were asked to rate the apparent gloss of each image using a 6-point scale with endpoints labelled “1 = completely matte” and “6 = extremely glossy”.

#### Experiment 2: Manipulating gloss

Experiment 2 measured gloss rankings for gloss-modulated network-generated images. Eighty sets of five 128×128 pixel (2.9×2.9cm) PixelVAE-generated images were shown. The five images within each set were arrayed in a random order in a row in the centre of the screen. Participants were asked to use the keyboard number keys to sort the images into a row of initially empty boxes at the bottom of the screen, in order from least glossy to most glossy. Each set of stimuli was shown once, for a total of 80 trials, presented in a random order.

#### Experiments 3a and 3b: Patterns of gloss constancy

Experiment 3 measured pairwise gloss comparisons for probe sequences differing only in surface relief. Experiments 3a and 3b differed only in the specific sequences shown to participants. Both experiments consisted of 10 sequences of 7 images, within which all possible pairs were shown twice each, for a total of 420 trials. Images appeared at a resolution of 800×800 pixels (18.6×18.6cm), side by side on screen. Side of screen and trial order were randomised, with all sequences interleaved with one another. Participants were asked to report, using the left and right arrow keys, which surface appeared more glossy.

#### Experiment 4: Degrees of gloss constancy

Experiment 4 measured pairwise gloss comparisons for pairs of images differing in surface relief or illumination. Each of 40 pairs of images was shown 8 times, for a total of 360 trials. Images appeared at a resolution of 128×128 pixels (2.9×2.9cm), side by side, with side of screen and trial order randomised. Participants reported, using the left and right arrow keys, which surface appeared more glossy.

### Data analysis

All analyses of human and model data were performed in Python 2.7 or 3.6, using numpy v1.16.5 and/or scikit-learn v0.21.3 packages.

#### Representational similarity analysis

We measured average correlation distance (1 minus Pearson *r*) between the latent representations of all 10,000 images in the ***main training dataset***, grouped by whether they used the same (diagonal blocks) or different (off-diagonal blocks) gloss/lighting conditions, for each network training instance. Representational dissimilarity matrices (RDMs; Figure 2D-E) were created by averaging these values over training instances of each model. To visualise the similarity of model predictions (Figure 7B) we created a vector, for each model, of predicted gloss values for all 270 rendered images used as experimental stimuli (50 images from Experiment 1; 140 images from Experiment 3; and 80 images from Experiment 4), normalised into the range 0–1. Euclidean distance between these prediction vectors was calculated for all pairs of models.

#### Decoding world factors from models and deriving gloss predictions

For models with multidimensional feature spaces (i.e., all except histogram skewness), a linear support vector machine (SVM) was used to classify specular reflectance (high vs low) of rendered images from their representations within the model. SVMs were trained on a random sample of 7,500 of the main training dataset images, and tested on the remaining 2,500 to derive gloss classification accuracies (Figure 3, Figure 6A and Supplementary Figures 4A and 5A). To derive continuously-valued gloss predictions for experimental images, images were input to each model, and the signed distance of their representation from the model’s SVM decision boundary was measured. Positive values indicate “high-gloss” classifications and negative values indicate “low-gloss”, with absolute magnitude indicating the strength of evidence for that classification (Figure 3, Figure 4B-C, Figure 5D-E, Figure 7A, Figure 8B, Supplementary Figures 6B-D and 7B).

For unsupervised PixelVAE and supervised ResNet models, we also trained linear SVMs to perform a 6-way light-field classification, and fitted linear regressions to predict surface relief (Figure 3B). Classifiers and regressions were performed once using the full 10D latent space for each network, and again using each of the network’s individual latent dimensions (Figure 3B; see also Supplementary Figure 1C for visualisation of individual dimensions in one unsupervised network).

For the histogram skewness model, raw skew values were used as gloss predictors for the purposes of model comparison. Ground-truth gloss classification accuracy (Figure 6A) was defined as the accuracy using an optimal threshold to separate high-from low-specular reflectance images from one another on the basis of their skewness, fitting the threshold on 7,500 main training dataset images and testing on the remaining 2,500.

#### Deriving predicted 2AFC experimental data from models

Model predictions for Experiment 3 were derived by simulating responses of each model in the 2AFC task performed by humans. For each sequence of 7 images, the predicted gloss values of each possible pair of images were compared, and the image for which the model predicted higher gloss was selected. Predicted responses were summarised as ‘proportion selected as being glossier’ for each level in the sequence, as for human data (Figure 5B and Supplementary Figure 3).

#### Measuring highlight features

Coverage, sharpness and contrast of highlights (Figure 8) were measured using a MATLAB package developed by Schmid et al ^115^. Briefly: (1) Coverage is defined as the proportion of pixels with higher intensities in the full image than in the diffuse component image; (2) Sharpness is defined as the Local Phase Coherence ^116^ within the specular component image; (3) Contrast is defined as the sum of the RMS contrast across eight bandpass-filtered versions of the specular component image.

### Psychophysical data analysis

No participants or trials were excluded from analysis. In Experiment 1 both human ratings and model-predicted gloss values were normalised to the range 0-1 before comparing, individually for each participant or model instance. In Experiment 2, rankings were converted to average rank position of each image step for each participant, averaging within matte and glossy seed image sequences. In Experiments 3a and 3b, pairwise comparisons were converted to the proportion of times each image within each sequence was judged as having higher gloss, across pairings and repetitions, for each participant. In Experiment 4, pairwise comparisons were converted to the proportion of times ‘image A’ (arbitrarily labelled) was judged as having higher gloss than ‘image B,’ and this proportion subtracted from 0.5 (i.e. 0 indicates equal apparent gloss, and good constancy; deviations in either direction indicate deviations from constancy). Model predictions were obtained by subtracting the predicted gloss value of image B from that of image A, and scaling this gloss difference into the range −0.5 to 0.5, retaining the original zero point. For models with multiple training instances, model predictions were always derived, and performances calculated, for each individual training instance.

### Statistical Analysis

Statistical analyses were performed using the Pingouin^117^ package for Python (version 0.3.10). All tests were two-tailed. Confidence intervals reported for mean differences and correlations were calculated by bootstrapping the respective estimate 10,000 times. Data distributions were assumed to be normal but this was not formally tested. The gloss predictions of all models were fixed, with no free parameters in evaluation against human data, except for the analysis of the data in Figure 5E, where the performance of a logistic transform of unsupervised model predictions is also reported.

## Data availability

All human and model data are available on Zenodo at http://doi.org/10.5281/zenodo.4495586.

## Code availability

Custom analysis code that supports the findings of this study is available on Zenodo at http://doi.org/10.5281/zenodo.4495586.

## Acknowledgements

This work was funded by the Deutsche Forschungsgemeinschaft (DFG, German Research Foundation; project number 222641018–SFB/TRR 135 TP C1), by the European Research Council (ERC; Consolidator Award ‘SHAPE’–project number ERC-CoG-2015-682859) to R.W.F. and by an Alexander von Humboldt Postdoctoral Research Fellowship to K.R.S.. The funders had no role in study design, data collection and analysis, decision to publish or preparation of the manuscript. We would like to thank Alexandra Schmid and Katja Doerschner for sharing code to implement the highlight feature measurement model, and Karl Gegenfurtner and James Todd for comments on earlier versions of this manuscript.

## Author contributions

Conceptualization, K.R.S., R.W.F and B.L.A..; Methodology, K.R.S. and R.W.F.; Software, K.R.S.; Formal Analysis, K.R.S.; Investigation, K.R.S.; Resources, R.W.F.; Writing – Original Draft, K.R.S. and R.W.F.; Writing – Review & Editing, K.R.S., R.W.F. and B.L.A.; Visualization, K.R.S; Supervision, R.W.F.; Funding Acquisition, R.W.F. and K.R.S..

## Supplementary Methods and Results

### Pilot experiment to select latent dimensionality of unsupervised network models

We wanted to select a latent dimensionality for the PixelVAE model such that the network was forced to learn a highly compressed code, but still able to capture the majority of meaningful structure in the training images. To gauge where this dimensionality lay for our dataset, we trained five PixelVAE networks with latent dimensionalities of 2, 5, 10, 20 and 40. From each network we generated 50 images by conditioning the generative sampling process on each of 50 random bumpy surface images that did not belong to the training dataset (see Supplementary Figure 1A). We then performed a psychophysical pilot experiment in which we showed twenty observers the generated images. On a trial, one of the rendered seed images was shown at the top of the screen, and its corresponding five generated images (one from each network) were shown in a random order in a horizontal row beneath. Participants rearranged the five generated images in order of their apparent similarity to the rendered seed image, and we measured the average similarity ranking given to the images generated by each model. Results are shown in Supplementary Figure 1B. Apparent similarity of generated images to seed images increased with a model’s latent dimensionality, but began to plateau after a dimensionality of ten, which we used for all subsequent PixelVAE models.

### Performance of intermediate layers in models

The analyses in the main work focus only on one 10-dimensional layer within both the supervised and unsupervised models. We also explored how well representations in the intermediate layers could predict both ground-truth gloss and human gloss perception, for one training instance of each model type. From the unsupervised PixelVAE architecture we trained an SVM to categorise high-vs-low gloss surfaces from the activations in each of the three convolutional layers of the autoencoder stream leading up to the latent code. From the supervised Resnet architecture we trained an SVM to categorise gloss from activations in the last convolutional layer of each each of the three residual blocks leading up to the output classification. Gloss classification accuracy was near-perfect for all layers of the supervised network, but increased across layers in the unsupervised network. Using the trained gloss classifiers, we then calculated predicted gloss values for each layer of each model and compared them to human gloss judgements in three psychophysical experiments as for all models in the main manuscript. The unsupervised network better predicted human gloss judgements across all layers, gradually improving across convolutional layers towards the 10-dimensional latent code (see Supplementary Figure 4A).

### Continuous gloss regression supervised network model

To explore whether a supervised network with a richer training objective might learn representations more similar to humans’, we created a version of the ResNet supervised model architecture in which the final two “high-vs-low-gloss” categorical units were replaced with a single continuously-valued “gloss level” unit (see Supplementary Figure 4B). A new training set of 10,000 images was rendered, in which the magnitude of the specular component was sampled randomly uniformly between 0 and 1, and the concentration of the specular component was set to the same value. All other scene factors were varied as in the original training set. This created a continuously-sampled one-dimensional gloss space, ranging from completely matte to completely mirrored. The supervised regression model was trained for 21 epochs on 9,000 of the images to predict the true specular magnitude, by minimising mean absolute error (MAE) using the Adam optimiser with a learning rate of 0.0001 and a decay rate of 0.9999 after each epoch. Other network parameters were kept as for the categorisation-supervised models. Five model instances were trained from different random initial weight settings. Ability to predict specular magnitude was excellent (average MAE on 500 test images = 0.002). We then evaluated the model’s ability to predict human data in the same way as for the categorisation-supervised ResNet models in the main manuscript, by training a gloss-classification SVM on the feature space in the 10D penultimate layer of the network, deriving gloss predictions for experimental images, and comparing them to human gloss judgements. (see Supplementary Figure 4B). The regression-supervised networks performed better than the categorisation-supervised networks, predicting human judgements on average as well as ground-truth specular reflectance (one-sample t-test of model RMSE against ground truth RMSE across model training instances *t_4_* = 0.24, *p* = 0.82, Cohen’s d = 0.10, 95% CI of difference = [-0.02–0.02]). Unsupervised models, however, consistently outperform ground-truth, predicting *errors* in human gloss perception, as well as successes (one-sample t-test of the poorest-performing set of unsupervised networks—those trained on a continuously-varying gloss dataset**—**against ground truth *t_4_* = −4.78, *p* = 0.009, d = 2.14, 95% CI = [0.02–0.08]; independent-samples t-test against regression-supervised networks *t_8_* = 3.98, *p* = 0.005, d = 2.52, 95% CI = [0.02–0.08]).

### Effect of varying model hyperparameters

When working with deep neural networks one must choose values for a large number of hyperparameters, such as the number of layers and convolutional filters, the learning rate, the rate at which the learning rate decays as training progresses, and more. Fully exploring this hyperparameter space is prohibitively time consuming, so we performed a small exploration of some key factors by training 28 additional models (14 unsupervised PixelVAE and 14 supervised ResNet networks) and evaluating their ability to classify gloss and to predict human gloss perception (see Supplementary Figure 5). Supplementary Table 1 lists the variants tested. For both the unsupervised and supervised models we tested a deeper and shallower architecture than the one originally used, as well as lower and higher learning rates, and lower and higher decay rates for the learning rate. Some hyperparameters of interest could not be varied in both model types. For batch size, we explored higher and lower sizes for the supervised model only, as memory constraints on the training GPUs limited the unsupervised model to the small batch size of 5. Finally, we explored higher and lower degrees of complexity for the pixel distribution learned by the unsupervised PixelVAE model, a hyperparameter not present in the supervised model. For each variant, we altered only the single hyperparameter of interest, and held all others at their original values. All models were trained for the same duration as the original models (200 epochs for PixelVAE models and 25 epochs for ResNet models, which was sufficient for convergence in each model type, see Supplementary Figure 5B). Once trained, each model instance was evaluated against human psychophysical data from Experiments 1, 3 and 4 identically to the model evaluation procedures in the main manuscript. Average error in predicting human data, across the three experiments, is shown for each network in Supplementary Figure 5A and C.

Unsupervised models generally predicted human perceptual data better than supervised models across the wide range of hyperparameter settings explored, with the exception of two unsupervised models with large learning rates that failed to train (as evidenced by poor convergence and noisy generated samples). The performances of the original unsupervised model training instances fall towards the better end of the range of performances found for hyperparameter variants, and those of the original supervised instances fall towards the worse end of the range of performances found for variants, but both are within the bounds apparently typical of their model types. The slightly larger difference between the original models may be due to having specifically selected stimuli for Experiments 3 and 4 in order to maximise the “disagreement” in gloss predictions made by the original models.

**Supplementary Table 1:** Hyperparameter values for each of 28 alternative variant models. For each variant, only the specified hyperparameter was altered, and all others were kept at the values used in the original models. The learning rate decay factor was used to gradually reduce the learning rate during training by multiplying the learning rate after each epoch by *1 − decay factor* (*learning rate* /*total epochs*). The PixelVAE model a pixel probability distribution over RGB space, which is parameterised as a mixture of logistic functions. The number of logistic functions is a hyperparameter which controls the complexity of the learned distribution.

| Hyperparameter | Values tested in unsupervised PixelVAE models | Values tested in supervised ResNet models | Value in original unsupervised PixelVAE model | Value in original supervised ResNet model |
| --- | --- | --- | --- | --- |
| Depth of network | 2 three-layer blocks, with 80 convolutional feature maps per layer<br>4 three-layer blocks, with 48 convolutional feature maps per layer | 2 three-layer blocks, with 80 convolutional feature maps per layer<br>4 three-layer blocks, with 48 convolutional feature maps per layer | 3 three-layer blocks, with 64 convolutional feature maps per layer<br>(in encoder part of PixelCNN stream) | 3 three-layer blocks, with 56 convolutional feature maps per layer |
| Learning rate | 0.01<br>0.005<br>0.0005<br>0.0001 | 0.01<br>0.005<br>0.0005<br>0.0001 | 0.001 | 0.001 |
| Decay factor for learning rate | 0.25<br>0.50<br>2.00<br>4.00 | 0.25<br>0.50<br>2.00<br>4.00 | 1.0 | 1.0 |
| Batch size |  | 12<br>22<br>42<br>50 | 5 (at memory capacity) | 32 |
| Complexity of PixelVAE's learned pixel distribution | 4 logistic functions<br>6 logistic functions<br>24 logistic functions<br>30 logistic functions |  | 12 logistic functions | n/a |

### Tolerance to variations in rendering method

In order to generate large datasets in a reasonable timeframe, we used real-time rasterised rendering in the Unity3D engine. The resulting images have good visual quality, but are not perfectly physically faithful. For example, they lack inter-reflections, and use an approximation of ambient occlusion. To test whether the unsupervised models trained on these rasterised images could make reasonable gloss predictions for surfaces rendered with more time-intensive but physically accurate methods, we generated 24 images of surfaces rendered once using rasterisation, and once physically faithful ray-tracing, via the Eevee and Cycles renders within the Blender engine, respectively. Each image was then input to all training instances of the original unsupervised PixelVAE model, and an average predicted gloss value was calculated. The models’ gloss classifications were correct for all images rendered via both methods, although gloss values were systematically slightly higher for images rendered via the rasterised method familiar to the networks (see Supplementary Figure 6B).

### Generalisation to real-world photographs

We performed two tests of generalisation to real-world images. First, we took 20 colour close-up photographs of common surfaces, ten of which were highly glossy (e.g. tomatoes, plastic, metallic foil) and ten of which were strongly matte (e.g. chalk, fleece, styrofoam). Predicted gloss values for each image were calculated for each of the ten training instances of the original unsupervised PixelVAE models (Supplementary Figure 6C). The unsupervised models were able to correctly classify all but one of the surfaces as being high or low gloss. Second, we performed a broader test by inputting to the models all images from the Giessen Material Image Database (Wiebel, Valsecchi & Gegenfurtner, 2013), which comprises 300 close-up photographs of assorted wood, metal, stone, and fabric surfaces. Predicted gloss values for each image from each model training instance are shown in Supplementary Figure 6D. Although the unsupervised model was moderately successful at sorting the “metal” category images from dull brushed surfaces to highly polished ones, it also showed clear failure cases. For example, matte fabrics with high-contrast patterns often received high predicted gloss values. Given that textured or patterned surfaces never occurred in the model’s training dataset, we would not expect it to have been able to learn the image structures associated with these.

## Supplementary Figures

**Supplementary Figure 1.**
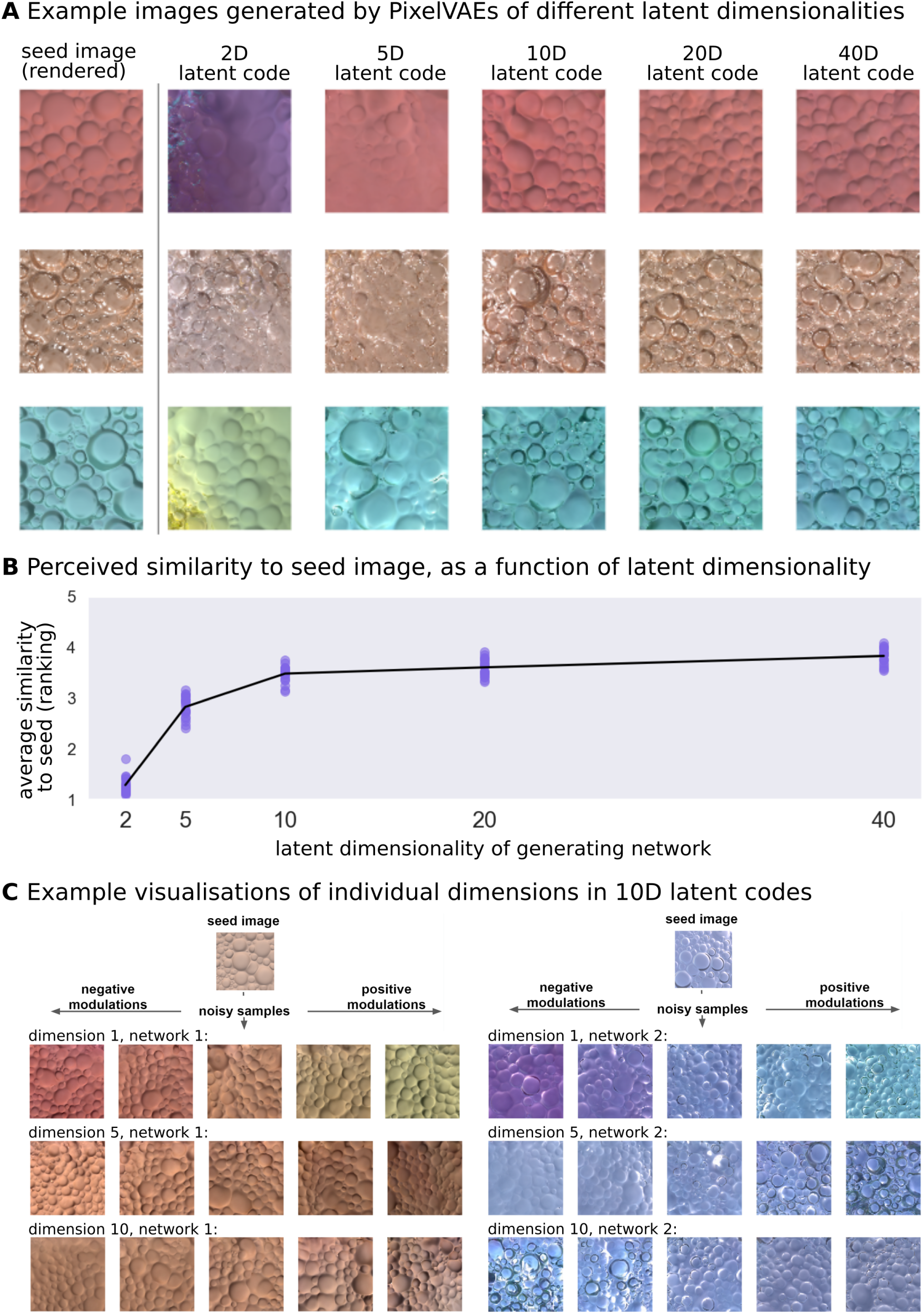
Choice of latent dimensionality, and visualisation of single dimensions. **(A)** Examples of images generated by networks with latent layers containing 2, 5, 10, 20, and 40 dimensions, using each of the three images in the leftmost column as “seeds” to select a point to sample in the latent space. Sample quality improves as latent dimensionality increases, up to a point. **(B)** Mean ranking of samples (y-axis) from each network (x-axis) in terms of perceived similarity to seed images, for each observer (coloured dots) in a psychophysical pilot experiment. Line indicates mean over all observers. Above around ten latent dimensions, there are diminishing perceptual quality returns when adding further dimensions, for this image training set. **(C)** Example visualisations of three individual dimensions within two different training instances of models with 10 latent dimensions, for two seed images. The central column beneath each seed image shows different samples generated by the network when sampled on the identical point in latent space. To generate the images in the two columns to the left of this, we conditioned on two points slightly negatively shifted, along only the stated dimension; to generate the images in the two right columns, we conditioned on two points slightly positively shifted along the same dimension. Individual dimensions capture variations in shape, colour, material, and lighting, and often combine multiple of these properties (i.e. the representation of world factors is distributed, not sparsely encoded by single dimensions).

**Supplementary Figure 2.**
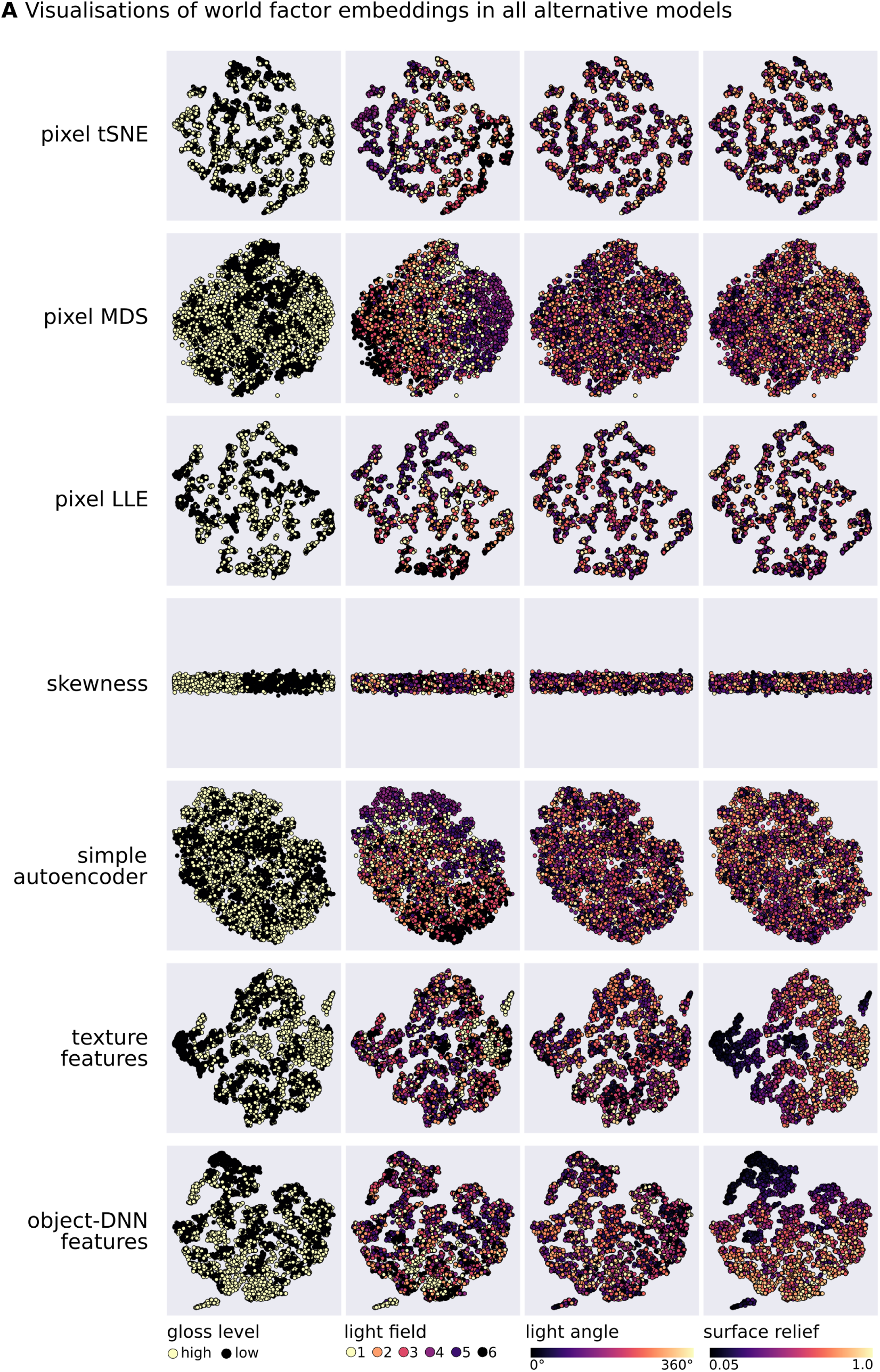
Visualisations of how world factors are embedded within the feature spaces of all alternative models. **(A)** Shown are visualisations of 4,000 random images from the training dataset, projected into two dimensions via t-weighted stochastic neighbourhood embedding for all models not shown in Figure 2 of the main manuscript. For each model, the 2D embedding is coloured by (from left to right) gloss level of the surface, light field illuminating the scene, rotation of light field with respect to the surface, and depth of surface relief. The luminance histogram skewness model (fourth row) is a one-dimensional feature space; some random vertical jitter has been added for easier visualisation. Most world factors are thoroughly intermingled in most of these alternative models, with a few exceptions. High and low gloss images are reasonably well separated by the simple statistic of luminance histogram skewness, within this dataset of cleanly bimodal gloss. Illumination fields are roughly separated within the pixel MDS and simple autoencoder feature spaces, likely because they influence the colour distributions in images. Interestingly, surface relief is best represented in the models consisting of texture features or features from a DNN trained to recognise objects in natural images, which also predict human judgements best among the alternative models.

**Supplementary Figure 3:**
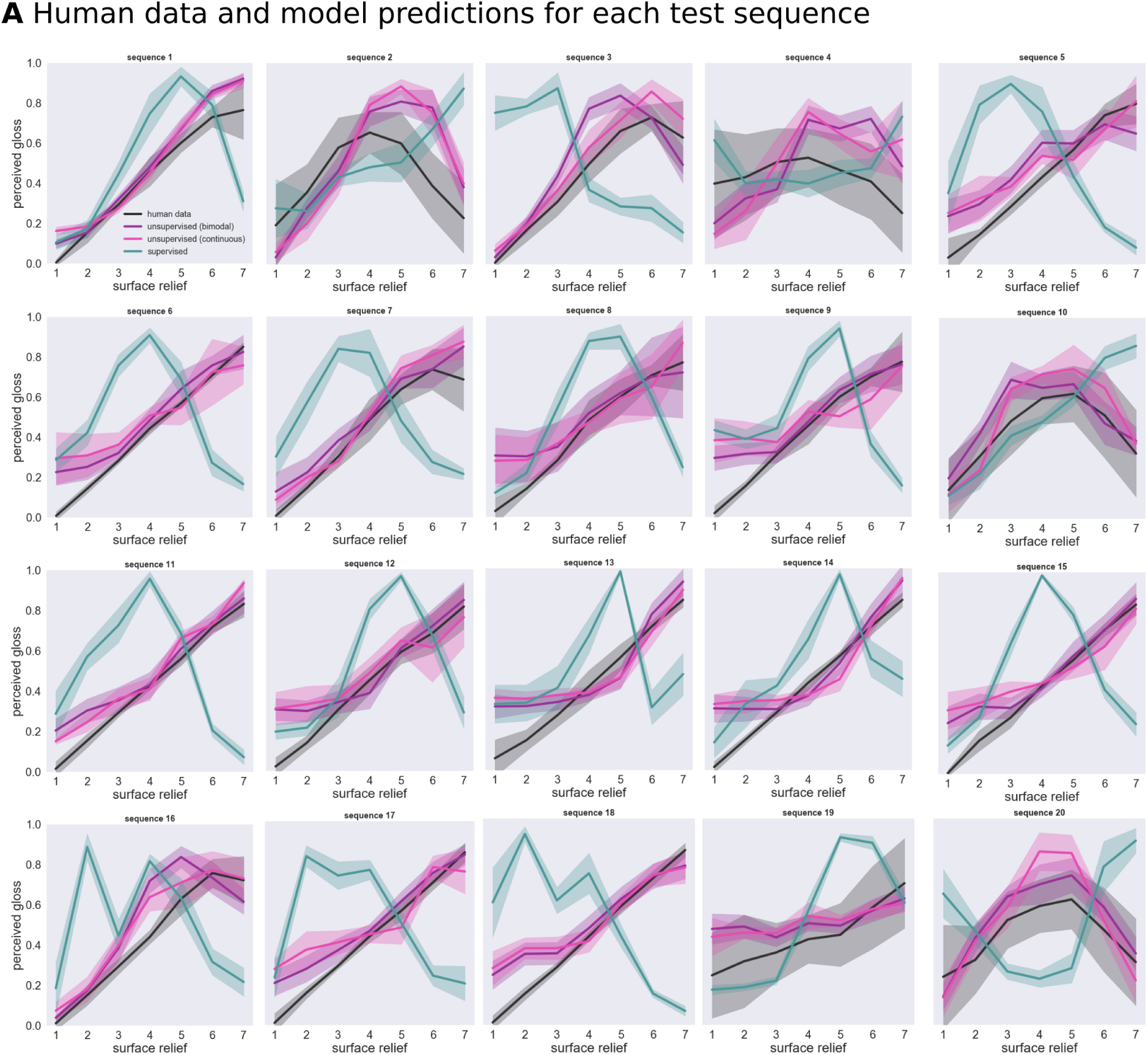
Full data for Experiment 3. **(A)** Full data from each of the twenty constancy sequences summarised in Figure 5B-C of the main text, including predictions from a new set of five PixelVAE models trained on an independent dataset in which gloss varies continuously rather than bimodally (“PixelVAE continuous-gloss” model referred to in main text and figures).

**Supplementary Figure 4:**
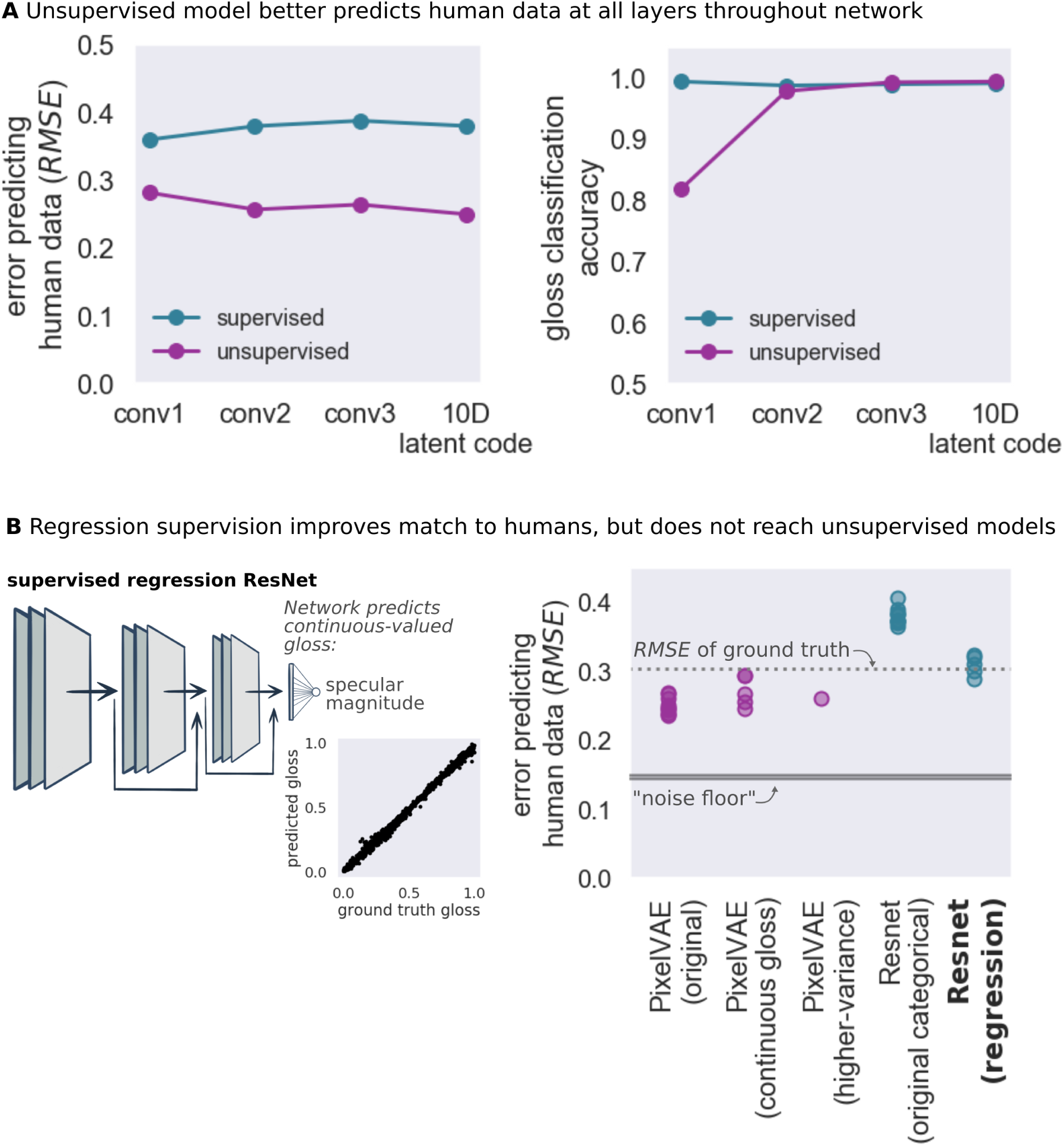
Main results are robust to changes in layer chosen for readout or training objective of supervised network. **(A)** Average error in predicting human gloss judgements across three psychophysical experiments (left) and objective gloss-classification accuracy (right) for successive convolutional layers of the unsupervised (purple) and supervised (teal) models, for one model training instance of each. The rightmost data points in each plot show the performance of the 10-dimensional latent code layer that forms the basis of all other analyses. The unsupervised model consistently better predicts human data across all its layers, whereas the supervised model displays near-perfect objective gloss classification throughout its layers. **(B)** Schematic (left) of the ResNet architecture adapted to output a single scalar value. After supervised training to predict specular magnitude, the model approximates ground-truth well (inset scatterplot shows true and predicted gloss levels for 500 unseen test images for one training instance of the network). Plot (right) shows average error in predicting human gloss judgements, across three psychophysical experiments, for the five training instances of the regression-supervised ResNet model (rightmost column). For comparison, performance is also shown for each training instance of (from left to right) the original unsupervised model, an unsupervised model trained on a dataset in which specular magnitude and concentration vary randomly and independently, an unsupervised model trained on a dataset in which surface geometry and illumination vary more widely, and for the original categorisation-supervised Resnet model. For reference, the dotted grey line indicates how well human judgements can be predicted by ground truth specular reflectance, and the shaded grey bar indicates how well human judgements can be predicted by data from other humans (the “noise floor”, or lowest possible model error).

**Supplementary Figure 5:**
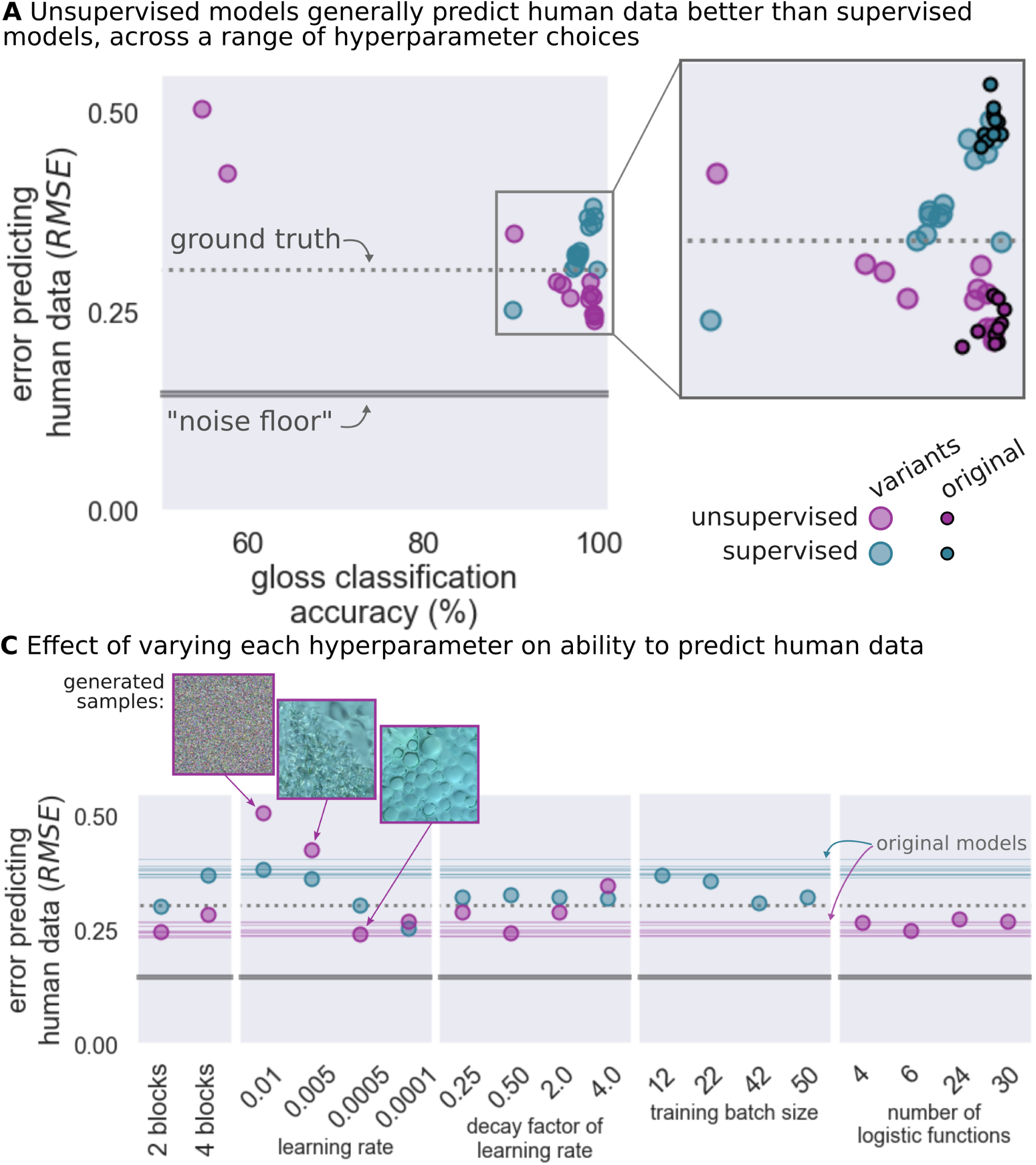
Main results are robust to hyperparameters of networks. **(A)** Scatterplot showing (y-axis) error in predicting human gloss judgements (average RMSE across three psychophysical experiments) against (x-axis) accuracy of high-vs-low gloss classifier trained on each network’s 10D latent code, for all 14 hyperparameter variants of the (purple) unsupervised PixelVAE and (teal) supervised Resnet model. Inset shows a magnified version of the region in which most models lie, excluding variants that failed to train well. Smaller black-outlined dots show the range of performances for the 10 original training instances of both model types. The dotted grey line indicates how well human judgements can be predicted by ground truth specular reflectance, and the shaded grey bar indicates how well human judgements can be predicted by data from other humans (the “noise floor”, or lowest possible model error). **(B)** Performance in predicting human gloss judgements (y-axis) for each of the 28 hyperparameter variant networks. Unsupervised networks were generally superior to supervised networks, with the exception of the two unsupervised networks with large learning rates which failed to converge during training. Inset images are samples generated from each of the PixelVAE networks with learning rates of 0.01, 0.005, and 0.0005, showing that at the largest learning rates, the model fails to learn the structure in the data. Models that were able to train, generally well predicted human perceptual judgements. The noise floor and ground-truth performance are shown as in A. Faint horizontal lines indicate the performance of each of the ten training instances of the original unsupervised (purple) and supervised (teal) models, giving an indication of the expected variation in performance due to differences in random initialisation.

**Supplementary Figure 6.**
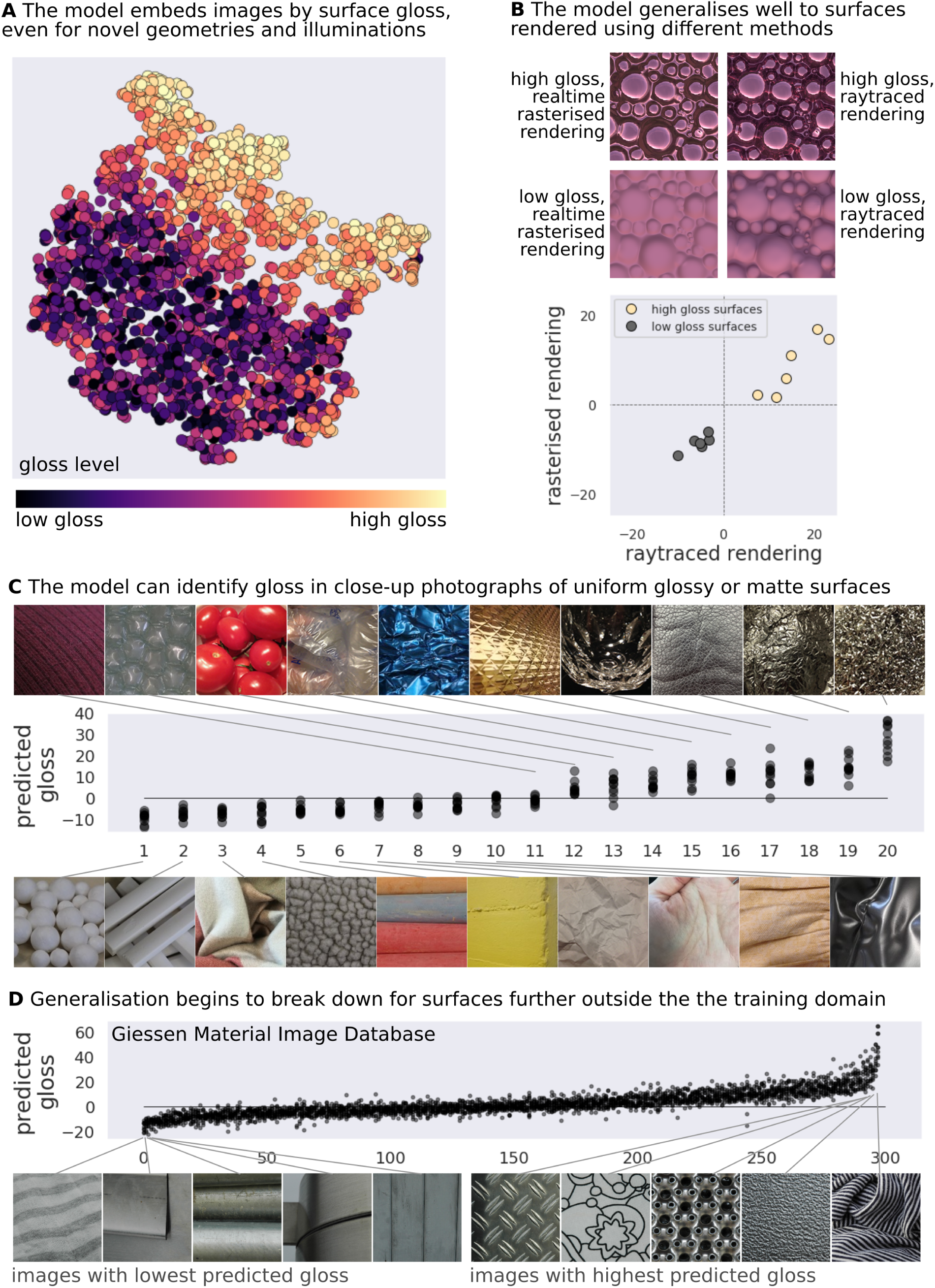
Generalisation to images outside the training set. **(A)** The trained networks organise images lawfully by material even for renderings that fall outside the parameters of the training dataset. Plot shows tSNE embedding of 2500 images generated using five surface geometries and eight lighting environments never sampled from within the network’s training dataset (see Methods: *Geometry and lighting generalisation test set)*. Points are colour coded by the strength of specular reflectance (dark purple = matte; cream = highest gloss). **(B)** Example images (top) of the same surface geometry with either high or low gloss, created using either approximate rasterised rendering (left two images) as in the experimental training and testing datasets, or physics-simulating raytraced rendering (right two images). Scatterplot (bottom) of gloss level predicted by the unsupervised PixelVAE models for each of 12 such surfaces when rendered using rasterisation as in the original experiments (x-axis) or ray-tracing (y-axis). Gloss values are closely correlated, and all images were correctly classified (i.e. high-gloss images receive positive gloss values, and low-gloss images receive negative values), regardless of rendering method. **(C)** Predicted gloss (y-axis) for twenty photographs of real-world glossy or matte surfaces, sorted by average predicted gloss. Each dot shows the prediction of one training instance of the unsupervised model. Different training instances make similar gloss predictions, and all but one surface is correctly categorised (i.e. matte surfaces receive negative gloss values (bottom row) and glossy surfaces receive positive values (top row)). **(D)** Predicted gloss for all 300 images of assorted wood, fabric, metal and stone surfaces comprising the Giessen Material Image Database (https://www.allpsych.uni-giessen.de/MID/), sorted by average predicted gloss. Bottom row shows the five images with the lowest (left) and highest (right) predicted gloss. Although the model correctly identifies gloss in e.g. polished metal surfaces, it also incorrectly predicts high gloss values for surfaces with high-contrast textures, such as patterned fabrics.

**Supplementary Figure 7.**
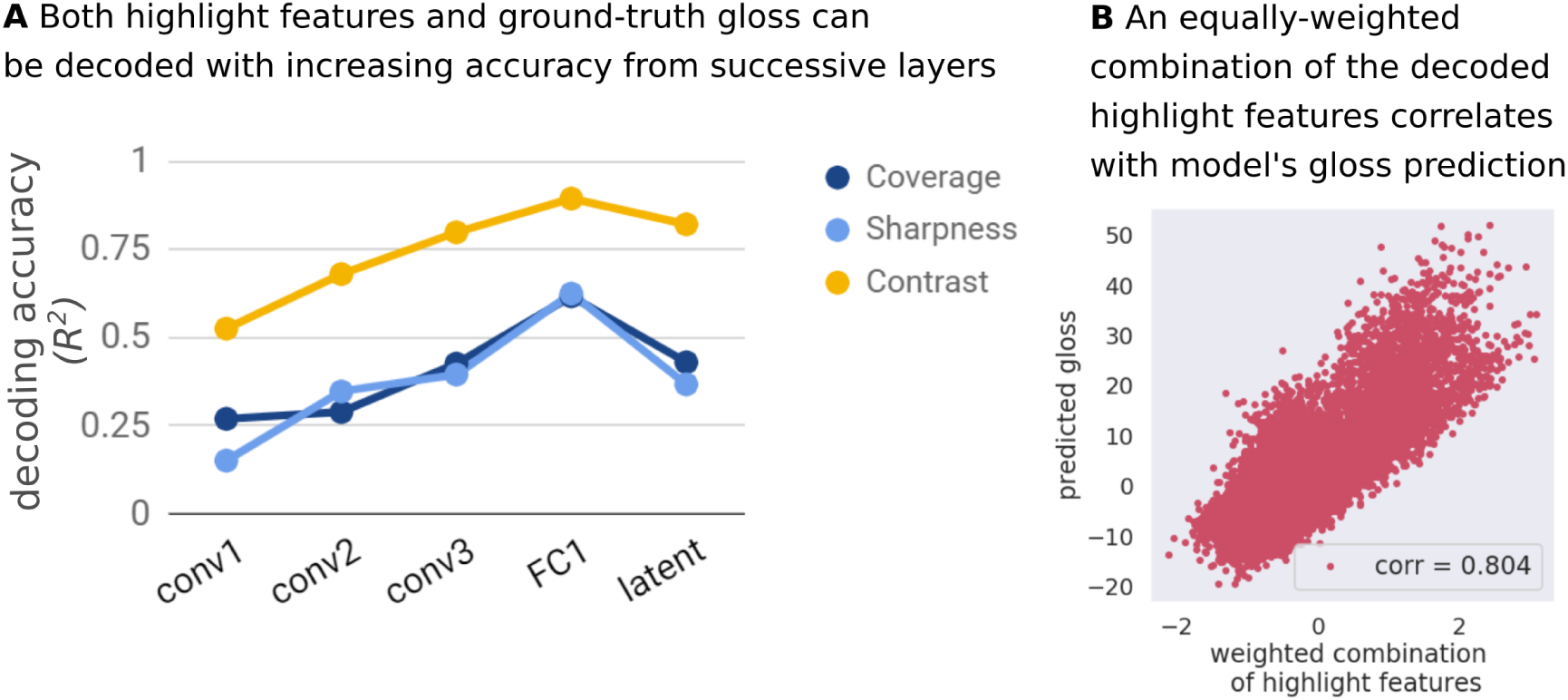
Highlight coverage, sharpness and contrast correlate with model-predicted gloss level. **(A)** The coverage, sharpness and contrast of highlights can be decoded from representations within one of the trained networks (here, we chose to investigate the model training instance with highest correlation with human-perceived gloss in Experiment 1). The decodability of mid-level highlight features increases throughout the network’s layers, until the bottleneck latent code, in which there is a slight decline. (However, note that the dimensionality of the feature space reduces from 1000 units in the final fully connected layer to only 10 in the latent layer, while only minimally affecting gloss classification performance). **(B)** Combining (via equal weighting) the estimates of the three highlight features, as decoded from the fully-connected layer immediately preceding the latent layer, yields a summary highlight measure which correlates well with the gloss level predicted from the latent layer.

## References

1. Adelson, E. H. Lightness Perception and Lightness Illusions. in The New Cognitive Neurosciences 339–351 (MIT Press, 2000).

2. Anderson, B. L. Mid-level vision. Current Biology 30, R105–R109 (2020).

3. Anderson, B. L. The perceptual representation of transparency, lightness, and gloss. Handbook of Perceptual Organization 466–483 (2015).

4. Barrow, H., Tenenbaum, J., Hanson, A. & Riseman, E. Recovering intrinsic scene characteristics. Computer Vision Systems 2, 2 (1978).

5. Fleming, R. W. Material perception. Annual Review of Vision Science 3, 365–388 (2017).

6. Todd, J. T. The visual perception of 3D shape. Trends in Cognitive Sciences 8, 115–121 (2004).

7. Todd, J. T., Norman, J. F. & Mingolla, E. Lightness constancy in the presence of specular highlights. Psychological Science 15, 33–39 (2004).

8. Marr, D. Vision. (Freeman, 1982).

9. Kersten, D., Mamassian, P. & Yuille, A. Object perception as Bayesian inference. Annual Review of Psychology 55, 271–304 (2004).

10. Geisler, W. S. & Kersten, D. Illusions, perception and Bayes. Nature Neuroscience 5, 508 (2002).

11. von Helmholtz, H. Treatise on physiological optics vol. iii. (1867).

12. Anderson, B. L. Can computational goals inform theories of vision? Topics in Cognitive Science 7, 274–286 (2015).

13. Hoffman, D. D., Singh, M. & Prakash, C. The interface theory of perception. Psychonomic Bulletin & Review 22, 1480–1506 (2015).

14. Fleming, R. W. & Storrs, K. R. Learning to see stuff. Current Opinion in Behavioral Sciences 30, 100–108 (2019).

15. Barlow, H. The exploitation of regularities in the environment by the brain. Behavioral and Brain Sciences 24, 602–607 (2001).

16. DiCarlo, J. J. & Cox, D. D. Untangling invariant object recognition. Trends in Cognitive Sciences 11, 333–341 (2007).

17. Storrs, K. R. & Fleming, R. W. Learning about the world by learning about images. Current Directions in Psychological Science (in press).

18. Higgins, I. et al. Towards a definition of disentangled representations. arXiv preprint arXiv:1812.02230 (2018).

19. Barlow, H. B. Possible principles underlying the transformation of sensory messages. Sensory Communication 1, 217–234 (1961).

20. Attneave, F. Some informational aspects of visual perception. Psychological Review 61, 183 (1954).

21. Simoncelli, E. P. & Olshausen, B. A. Natural image statistics and neural representation. Annual Review of Neuroscience 24, 1193–1216 (2001).

22. Olshausen, B. A. & Field, D. J. Emergence of simple-cell receptive field properties by learning a sparse code for natural images. Nature 381, 607–609 (1996).

23. Grossberg, S. Adaptive pattern classification and universal recoding: I. Parallel development and coding of neural feature detectors. Biological Cybernetics 23, 121–134 (1976).

24. Földiak, P. Forming sparse representations by local anti-Hebbian learning. Biological Cybernetics 64, 165–170 (1990).

25. Anderson, B. L. Visual perception of materials and surfaces. Current Biology 21, R978–R983 (2011).

26. Gilchrist, A. et al. An anchoring theory of lightness perception. Psychological Review 106, 795 (1999).

27. Pont, S. C. & te Pas, S. F. Material—Illumination ambiguities and the perception of solid objects. Perception 35, 1331–1350 (2006).

28. Adams, W. J., Kucukoglu, G., Landy, M. S. & Mantiuk, R. K. Naturally glossy: Gloss perception, illumination statistics, and tone mapping. Journal of Vision 18, 4–4 (2018).

29. Foster, D. H. Color constancy. Vision Research 51, 674–700 (2011).

30. Motoyoshi, I. & Matoba, H. Variability in constancy of the perceived surface reflectance across different illumination statistics. Vision Research 53, 30–39 (2012).

31. Chadwick, A. C. & Kentridge, R. The perception of gloss: A review. Vision Research 109, 221–235 (2015).

32. Obein, G., Knoblauch, K. & Viéot, F. Difference scaling of gloss: Nonlinearity, binocularity, and constancy. Journal of Vision 4, 4–4 (2004).

33. Fleming, R. W., Dror, R. O. & Adelson, E. H. Real-world illumination and the perception of surface reflectance properties. Journal of Vision 3, 3–3 (2003).

34. Ho, Y.-X., Landy, M. S. & Maloney, L. T. Conjoint measurement of gloss and surface texture. Psychological Science 19, 196–204 (2008).

35. Marlow, P. J., Kim, J. & Anderson, B. L. The Perception and Misperception of Specular Surface Reflectance. Current Biology 22, 1909–1913 (2012).

36. Doerschner, K. et al. Visual motion and the perception of surface material. Current Biology 21, 2010–2016 (2011).

37. Wendt, G., Faul, F., Ekroll, V. & Mausfeld, R. Disparity, motion, and color information improve gloss constancy performance. Journal of Vision 10, 7–7 (2010).

38. Toscani, M., Guarnera, D., Guarnera, C., Hardeberg, J. Y. & Gegenfurtner, K. Three perceptual dimensions for specular and diffuse reflection. ACM Transactions on Applied Perception (2020).

39. Ferwerda, J. A., Pellacini, F. & Greenberg, D. P. Psychophysically based model of surface gloss perception. in Human Vision and Electronic Imaging VI vol. 4299 291–301 (International Society for Optics and Photonics, 2001).

40. Lagunas, M. et al. A Similarity Measure for Material Appearance. ACM Transactions on Graphics (SIGGRAPH 2019) 38, (2019).

41. Ingersoll, L. R. The Glarimeteran Instrument for Measuring the Gloss of Paper. Journal of the Optical Society of America 5, 213–217 (1921).

42. Ward, G. J. Measuring and modeling anisotropic reflection. in Proceedings of the 19th annual conference on Computer Graphics and Interactive Techniques 265–272 (1992).

43. Wills, J., Agarwal, S., Kriegman, D. & Belongie, S. Toward a perceptual space for gloss. ACM Transactions on Graphics (TOG) 28, 1–15 (2009).

44. Serrano, A., Gutierrez, D., Myszkowski, K., Seidel, H.-P. & Masia, B. An intuitive control space for material appearance. ACM Transactions on Graphics (SIGGRAPH ASIA 2016) 35, (2016).

45. Vangorp, P., Laurijssen, J. & Dutré, P. The influence of shape on the perception of material reflectance. in ACM SIGGRAPH 2007 77 (2007).

46. Salakhutdinov, R. Learning deep generative models. Annual Review of Statistics and Its Application 2, 361–385 (2015).

47. Zhao, S., Song, J. & Ermon, S. Towards deeper understanding of variational autoencoding models. arXiv preprint arXiv:1702.08658 (2017).

48. Gulrajani, I., et al. Pixelvae: A latent variable model for natural images. arXiv preprint arXiv:1611.05013 (2016).

49. Radford, A., Metz, L. & Chintala, S. Unsupervised representation learning with deep convolutional generative adversarial networks. arXiv preprint arXiv:1511.06434 (2015).

50. Higgins, I. et al. beta-VAE: Learning Basic Visual Concepts with a Constrained Variational Framework. International Conference on Learning Representations 2, 6 (2017).

51. Lindsay, G. Convolutional neural networks as a model of the visual system: past, present, and future. Journal of Cognitive Neuroscience 1–15 (2020).

52. Yamins, D. L. & DiCarlo, J. J. Using goal-driven deep learning models to understand sensory cortex. Nature Neuroscience 19, 356 (2016).

53. Storrs, K. R. & Kriegeskorte, N. Deep learning for cognitive neuroscience. in The Cognitive Neurosciences (MIT Press, 2020).

54. Richards, B. A. et al. A deep learning framework for neuroscience. Nature Neuroscience 22, 1761–1770 (2019).

55. Kriegeskorte, N. Deep neural networks: a new framework for modeling biological vision and brain information processing. Annual Review of Vision Science 1, 417–446 (2015).

56. LeCun, Y., Bengio, Y. & Hinton, G. Deep learning. Nature 521, 436–444 (2015).

57. He, K., Zhang, X., Ren, S. & Sun, J. Deep residual learning for image recognition. in Proceedings of the IEEE conference on Computer Vision and Pattern Recognition 770–778 (2016).

58. Russakovsky, O. et al. Imagenet large scale visual recognition challenge. International Journal of Computer Vision 115, 211–252 (2015).

59. Taigman, Y., Yang, M., Ranzato, M. & Wolf, L. Deepface: Closing the gap to human-level performance in face verification. in Proceedings of the IEEE conference on Computer Vision and Pattern Recognition 1701–1708 (2014).

60. Cadieu, C. F. et al. Deep neural networks rival the representation of primate IT cortex for core visual object recognition. PLoS Computational Biology 10, e1003963 (2014).

61. Schrimpf, M., et al. Brain-Score: Which artificial neural network for object recognition is most brain-like? bioRxiv preprint (2018).

62. Storrs, K. R., Kietzmann, T. C., Walther, A., Mehrer, J. & Kriegeskorte, N. Diverse deep neural networks all predict human IT well, after training and fitting. Journal of Cognitive Neuroscience (in press).

63. Khaligh-Razavi, S.-M. & Kriegeskorte, N. Deep supervised, but not unsupervised, models may explain IT cortical representation. PLoS Computational Biology 10, (2014).

64. Nguyen, A., Yosinski, J. & Clune, J. Deep neural networks are easily fooled: High confidence predictions for unrecognizable images. in Proceedings of the IEEE conference on Computer Vision and Pattern Recognition 427–436 (2015).

65. Geirhos, R. et al. Generalisation in humans and deep neural networks. in Advances in Neural Information Processing Systems 7538–7550 (2018).

66. Geirhos, R. et al. ImageNet-trained CNNs are biased towards texture; increasing shape bias improves accuracy and robustness. arXiv preprint arXiv:1811.12231 (2018).

67. Geirhos, R. et al. Shortcut Learning in Deep Neural Networks. arXiv preprint arXiv:2004.07780 (2020).

68. Kingma, D. P. & Welling, M. Auto-encoding variational bayes. arXiv preprint arXiv:1312.6114 (2013).

69. Hinton, G. E. & Salakhutdinov, R. R. Reducing the dimensionality of data with neural networks. Science 313, 504–507 (2006).

70. Mehrer, J., Spoerer, C. J., Kriegeskorte, N. & Kietzmann, T. C. Individual differences among deep neural network models. Nature Communications 11, (2020).

71. He, K., Zhang, X., Ren, S. & Sun, J. Delving deep into rectifiers: Surpassing human-level performance on imagenet classification. in Proceedings of the IEEE international conference on Computer Vision 1026–1034 (2015).

72. Bengio, Y., Courville, A. & Vincent, P. Representation learning: A review and new perspectives. IEEE Transactions on Pattern Analysis and Machine Intelligence 35, 1798–1828 (2013).

73. Testolin, A., Stoianov, I. & Zorzi, M. Letter perception emerges from unsupervised deep learning and recycling of natural image features. Nature Human Behaviour 1, 657–664 (2017).

74. Maaten, L. van der & Hinton, G. Visualizing data using t-SNE. Journal of Machine Learning Research 9, 2579–2605 (2008).

75. Roweis, S. T. & Saul, L. K. Nonlinear dimensionality reduction by locally linear embedding. Science 290, 2323–2326 (2000).

76. Nili, H. et al. A toolbox for representational similarity analysis. PLoS Computational Biology 10, (2014).

77. Kriegeskorte, N. & Kievit, R. A. Representational geometry: integrating cognition, computation, and the brain. Trends in Cognitive Sciences 17, 401–412 (2013).

78. Kriegeskorte, N. & Diedrichsen, J. Inferring brain-computational mechanisms with models of activity measurements. Philosophical Transactions of the Royal Society B: Biological Sciences 371, 20160278 (2016).

79. Testolin, A. & Zorzi, M. Probabilistic models and generative neural networks: Towards an unified framework for modeling normal and impaired neurocognitive functions. Frontiers in Computational Neuroscience 10, 73 (2016).

80. Hong, H., Yamins, D. L., Majaj, N. J. & DiCarlo, J. J. Explicit information for category-orthogonal object properties increases along the ventral stream. Nature Nneuroscience 19, 613 (2016).

81. Naselaris, T., Kay, K. N., Nishimoto, S. & Gallant, J. L. Encoding and decoding in fMRI. NeuroImage 56, 400–410 (2011).

82. Gatys, L., Ecker, A. S. & Bethge, M. Texture synthesis using convolutional neural networks. In Advances in Neural Information Processing Systems 262–270 (2015).

83. Zhang, R., Isola, P., Efros, A. A., Shechtman, E. & Wang, O. The unreasonable effectiveness of deep features as a perceptual metric. in Proceedings of the IEEE Conference on Computer Vision and Pattern Recognition 586–595 (2018).

84. Rajalingham, R. et al. Large-scale, high-resolution comparison of the core visual object recognition behavior of humans, monkeys, and state-of-the-art deep artificial neural networks. Journal of Neuroscience 38, 7255–7269 (2018).

85. Portilla, J. & Simoncelli, E. P. A parametric texture model based on joint statistics of complex wavelet coefficients. International Journal of Computer Vision 40, 49–70 (2000).

86. Motoyoshi, I., Nishida, S., Sharan, L. & Adelson, E. H. Image statistics and the perception of surface qualities. Nature 447, 206–209 (2007).

87. Funke, C. M. et al. The Notorious Difficulty of Comparing Human and Machine Perception. arXiv preprint arXiv:2004.09406 (2020).

88. Golan, T., Raju, P. C. & Kriegeskorte, N. Controversial stimuli: pitting neural networks against each other as models of human recognition. arXiv preprint arXiv:1911.09288 (2019).

89. Wang, Z. & Simoncelli, E. P. Maximum differentiation (MAD) competition: A methodology for comparing computational models of perceptual quantities. Journal of Vision 8, 8–8 (2008).

90. Havran, V., Filip, J. & Myszkowski, K. Perceptually motivated BRDF comparison using single image. in Computer Graphics Forum vol. 35 1–12 (Wiley Online Library, 2016).

91. Wiebel, C. B., Valsecchi, M. & Gegenfurtner, K. R. The speed and accuracy of material recognition in natural images. Attention, Perception, & Psychophysics 75, 954–966 (2013).

92. Beck, J. & Prazdny, S. Highlights and the perception of glossiness. Perception & Psychophysics (1981).

93. Anderson, B. L. & Kim, J. Image statistics do not explain the perception of gloss and lightness. Journal of Vision 9, 10–10 (2009).

94. Marlow, P. J., Todorović, D. & Anderson, B. L. Coupled computations of three-dimensional shape and material. Current Biology 25, R221–R222 (2015).

95. Marlow, P. J. & Anderson, B. L. Material properties derived from three-dimensional shape representations. Vision Research 115, 199–208 (2015).

96. Marlow, P. J. & Anderson, B. L. Generative constraints on image cues for perceived gloss. Journal of Vision 13, 2–2 (2013).

97. Simoncelli, E. P. Vision and the statistics of the visual environment. Current Opinion in Neurobiology 13, 144–149 (2003).

98. Sawayama, M. & Nishida, S. Material and shape perception based on two types of intensity gradient information. PLoS Computational Biology 14, e1006061 (2018).

99. Nishida, S. & Shinya, M. Use of image-based information in judgments of surface-reflectance properties. Journal of the Optical Society of America A 15, 2951–2965 (1998).

100. Adelson, E. H. & Pentland, A. P. The perception of shading and reflectance. Perception as Bayesian Inference 409–423 (1996).

101. Marlow, P. J. & Anderson, B. L. Motion and texture shape cues modulate perceived material properties. Journal of Vision 16, 5–5 (2016).

102. Wiesel, T. N. & Hubel, D. H. Ordered arrangement of orientation columns in monkeys lacking visual experience. Journal of Comparative Neurology 158, 307–318 (1974).

103. Yang, J., Otsuka, Y., Kanazawa, S., Yamaguchi, M. K. & Motoyoshi, I. Perception of surface glossiness by infants aged 5 to 8 months. Perception 40, 1491–1502 (2011).

104. Balas, B. Children’s use of visual summary statistics for material categorization. Journal of Vision 17, 22–22 (2017).

105. Balas, B., Auen, A., Thrash, J. & Lammers, S. Children’s use of local and global visual features for material perception. Journal of Vision 20, 10–10 (2020).

106. Smith, L. B. & Slone, L. K. A developmental approach to machine learning? Frontiers in Psychology 8, 2124 (2017).

107. Pouget, A., Beck, J. M., Ma, W. J. & Latham, P. E. Probabilistic brains: knowns and unknowns. Nature Neuroscience 16, 1170 (2013).

108. Friston, K. The free-energy principle: a rough guide to the brain? Trends in Cognitive Sciences 13, 293–301 (2009).

109. Deneve, S. Bayesian spiking neurons I: inference. Neural Computation 20, 91–117 (2008).

110. Brainard, D. H. et al. Functional consequences of the relative numbers of L and M cones. Journal of the Optical Society of America A 17, 607–614 (2000).

111. Smirnakis, S. M., Berry, M. J., Warland, D. K., Bialek, W. & Meister, M. Adaptation of retinal processing to image contrast and spatial scale. Nature 386, 69–73 (1997).

112. Fleming, R. W. Visual perception of materials and their properties. Vision Research 94, 62–75 (2014).

113. Salimans, T., Karpathy, A., Chen, X. & Kingma, D. P. Pixelcnn++: Improving the pixelcnn with discretized logistic mixture likelihood and other modifications. International Conference on Learning Representations 2, (2017).

114. Van den Oord, A. et al. Conditional image generation with pixelcnn decoders. in Advances in Neural Information Processing Systems 4790–4798 (2016).

115. Schmid, A. C., Barla, P. & Doerschner, K. Material category determined by specular reflection structure mediates the processing of image features for perceived gloss. bioRxiv 2019–12 (2020).

116. Hassen, R., Wang, Z. & Salama, M. M. A. Image Sharpness Assessment Based on Local Phase Coherence. IEEE Transactions on Image Processing 22, 2798–2810 (2013).

117. Vallat, R. Pingouin: statistics in Python. Journal of Open Source Software 3, 1026 (2018).

